# seekrflow: Towards end-to-end automated simulation pipeline with machine-learned force fields for accelerated drug-target kinetic and thermodynamic predictions

**DOI:** 10.1101/2025.08.13.669965

**Authors:** Anupam A. Ojha, Lane W. Votapka, Shiksha Dutta, Anson F. Noland, Sonya M. Hanson, Rommie E. Amaro

## Abstract

Accurate prediction of drug-target binding and unbinding kinetics and thermodynamics is essential for guiding drug discovery and lead optimization. However, traditional atomistic simulations are often too computationally expensive to capture rare events that govern ligand (un)binding. Several enhanced sampling methods exist to overcome these limitations, but they require extensive manual intervention and introduce variability and artifacts in free energy and kinetic estimates that limit high-throughput scalability. The present work introduces seekrflow, an automated multiscale milestoning simulation pipeline that streamlines the entire workflow from a single receptor-ligand input structure to kinetic and thermodynamic predictions in a single step. This integrated approach minimizes manual intervention, reduces computational overhead, and enhances the reproducibility and accuracy of kinetic and thermodynamic predictions. The accuracy and efficiency of the pipeline is demonstrated on multiple receptor-ligand complexes, including inhibitors of heat shock protein 90, threonine-tyrosine kinase, and the trypsin protein, with predicted kinetic parameters closely matching experimental estimates. seekrflow establishes a new benchmark for automated and high-throughput physics-based predictions of kinetics and thermodynamics.

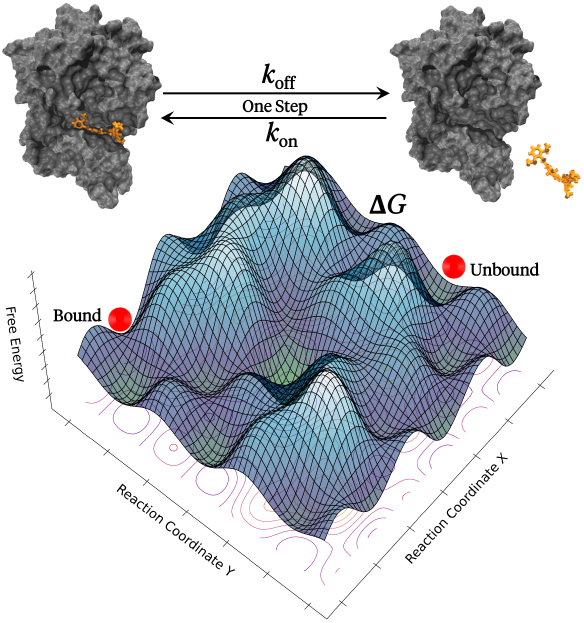

## 1 Introduction

Understanding receptor-ligand kinetics is essential for drug discovery applications, as these interactions govern fundamental processes such as molecular recognition, signal transduction, and enzymatic regulation ^1,2^. The rates at which an inhibitor binds to and dissociates from its receptor directly influence drug efficacy, determining how long a molecule remains active in its target site ^3–7^. Kinetic parameters such as *k*_on_ and *k*_off_ determine whether a ligand binds quickly and transiently or slowly and persistently. This distinction is particularly relevant in developing non-covalent inhibitors, allosteric modulators, and drugs targeting dynamic protein conformations ^8–10^.

The drug discovery community has long focused on improving free energy calculations for drug binding and unbinding, leading to high-affinity drug candidates ^11–13^. However, a high binding affinity determined by free energy calculations does not necessarily translate to a long residence time at its target ^14,15^, since kinetic factors (*k*_on_ and *k*_off_) dictate the duration of target engagement and therefore, the pharmocodynamic response. This is because drugs are only as potent as long as they are bound to their targets– numerous potent inhibitors with high affinity have failed in clinical trials due to short drug-target engagements ^16^. Drugs with slow dissociation rates have prolonged therapeutic effects, whereas those with fast dissociation rates may require higher or more frequent dosages to maintain efficacy. Kinetic parameters such as association (*k*_on_) and dissociation (*k*_off_) rates provide insights into drug selectivity by estimating binding duration across closely related receptors. This enables inhibitor design with minimized off-target effects and optimized pharmacodynamics. Studies have shown a strong correlation between prolonged pharmacological effects with drug-target residence times ^17,18^. A recent study developed a medium-throughput kinetic assay to measure for unlabelled human M_3_ receptor, demonstrating that a longer residence time drives *in vivo* efficacy ^17^. Similarly, another study developed a mechanistic pharmokinetics (PK)-pharmodynamics (PD) model based on prolonged drug-target residence times for LpxC inhibitors that translated into *in vivo* and cellular efficacy ^18^. It has also been demonstrated that long-lived drug-target complexes have yielded superior efficacy despite moderate affinity ^16,18–23^.

Despite such evidence, the significance of drug-target kinetics is still to be recognized to its full potential. Traditional drug discovery pipelines favor predicting equilibrium constants over estimating residence times. This is partly due to the existence of several standardized and user-friendly computational tools for affinity estimation. Atomistic molecular dynamics (MD) simulations, combined with free energy perturbation (FEP) methods such as Schrödinger’s FEP+ ^24^, AMBER thermodynamic integration (TI) ^25^, and approaches such as molecular mechanics generalized Born surface area (MM/GBSA) and molecular mechanics Poisson-Boltzmann surface area (MM/PBSA) ^26^, have become standard tools for estimating equilibrium binding affinity (ΔG), dissociation constant K_d_, or the inhibition constant K_i_ or IC_50_. Secondly, the prediction of kinetic parameters is often limited by the high computational cost of long-timescale MD simulations to achieve equilibrium and observe dissociation/association events ^27–31^. Recent computational methods estimate residence times more efficiently. However, issues related to reproducibility, automation, and manual intervention restrict their widespread use. In parallel, experimental techniques such as surface plasmonic resonance (SPR) ^32,33^ and stopped-flow measurements ^34^ remain low-throughput. We, therefore, require improved strategies for accurate and scalable kinetic predictions.

### 1.1 Sampling approaches for residence time predictions

Recent methods for residence time predictions (*k*_off_^−1^) can be broadly categorized into two main categories: (i) biased simulation approaches that correlate residence times with the escape times observed in biased MD simulations, such as random acceleration MD (RAMD) ^35,36^, Gaussian-accelerated MD (GaMD) ^37^, scaled MD ^38–40^, and metadynamics ^41^, and (ii) rigorous unbiased simulations that reconstruct the original timescales through statistical mechanics frameworks such as weighted ensemble ^42–44^, milestoning ^45–47^, and Markov state models ^48–50^. While biased simulations provide rapid insights into ligand escape pathways, these approaches, by definition, introduce artificial biases that may distort kinetic and thermodynamic estimates. On the other hand, unbiased simulations provide reliable and reproducible absolute residence time predictions but demand significant computational resources and complex setups. This limits their practicality for fast-paced lig- and optimization during early-stage drug discovery campaigns.

Given sufficient sampling, conventional MD simulations can quantify ligand (un)binding kinetics. Exceptionally long, unbiased MD simulations run on specialized hardware have been shown to capture ligand unbinding events with kinetic rates closer to experimental values ^51–53^. Such approaches are computationally expensive and impractical for routine use due to the enormous lengths of simulations required, often ranging from weeks to months. To accelerate such unbinding events, enhanced sampling approaches are therefore employed to pull the ligand out from the binding site by applying an external bias to the system with carefully chosen collective variables (CVs). Several enhanced sampling approaches (discussed next) have demonstrated some success in estimating the kinetics and thermodynamics of protein-ligand (un)binding.

Markov state models (MSMs) provide a framework to discretize MD simulation trajectories (or protein-ligand conformational space) into a network of metastable states to estimate transition probabilities and provide both kinetics and thermodynamics of protein-ligand (un)binding ^48–50^. For example, an MSM built on extensive unbiased MD simulation (often hundreds of microseconds) accurately predicted the association and dissociation rates of the trypsin-benzamidine complex ^54,55^. Once constructed, MSMs provide mechanistic insights into ligand unbinding pathways and large-scale protein conformational changes, aiding drug discovery processes. Building an MSM is expensive and non-trivial, often requiring extensive simulation over hundreds of microseconds. The lag time for the model has to be carefully tuned by selecting the shortest interval where the implied timescale is approximately constant ^56–58^. Errors in estimating kinetic rates often occur due to poor clustering of conformations that violate the Markov assumption or insufficient sampling of the ultra-stable complex. Compared to MSMs, weighted ensemble (WE) simulations is an alternative enhanced unbiased sampling scheme ^44^. WE is an unbiased sampling approach that employs an ensemble of trajectories with periodic reweighting, while pruning or merging trajectories to maintain a constant flux ^42,50^. The probabilistic redistribution of statistical weights of trajectories leads to rapid exploration of rare events. Once a steady state is reached, kinetic rate constants are measured using the Hill relation ^61^. WE is an unbiased and statistically exact sampling approach that captures ligand unbinding pathways and short and long-lived intermediates ^62^. However, an *a priori* definition of the CV is required where the phase space is partitioned into bins for resampling and reweighting purposes. A high computational cost associated with prolonged statistical convergence times makes WE a challenging approach to estimate kinetic rates for a series of drug-target complexes ^63,64^.

Metadynamics accelerates the sampling of protein-ligand (un)binding events by adding bias along the CVs, exploring the free energy landscapes inaccessible on unbiased MD simulation timescales ^41^. In infrequent metadynamics, the bias deposition is scarce, allowing the transition to occur naturally ^65,66^. True kinetics is then recovered by reweighting the waiting times using the cumulative bias introduced along the trajectory. This approach requires an *a priori* definition of appropriate CVs, which demands a detailed understanding of the drug-target complex. Suboptimal CVs may lead to unknown barriers and non-physical pathways. Complex free energy landscapes often require long biasing times, where errors of several kcal/mol are common. The height and the width of the Gaussian bias, along with its deposition frequency, must be carefully tuned to avoid overshooting transitions or getting stuck in artificial minima ^67^. Moreover, the height and shape of the true transition barrier is perturbed by the deposited bias, which alters the transition-state ensembles. Such perturbations can lead to distorted kinetics if the bias has significantly distorted the underlying dynamics of the ligand unbinding pathways.

Random acceleration MD (RaMD) employs a randomly oriented force on the ligand to pull it out of the binding site into the solvent ^35,68^. This adaptive force is periodically reassigned until the ligand moves beyond a certain distance threshold, allowing for a ligand dissociation on an accelerated timescale. One extension of this method is the *τ*-RaMD, which employs multiple unbinding trajectories to estimate relative residence times for drug-target complexes ^36^. RaMD/*τ*-RaMD do not require an *a priori* CV definition, often yielding multiple unbinding pathways and metastable states. Relative residence times for a series of inhibitors can therefore be achieved with certain accuracy, but true kinetics are difficult to achieve due to non-physical perturbation of the free energy landscape by the applied random force that distorts the barrier-crossing statistics. Similar to RaMD/*τ*-RaMD, Gaussian accelerated MD (GaMD) accelerates sampling of protein-ligand unbinding by applying a harmonic boost to the potential energy of the system ^69–72^. Smoothening of the free energy landscape lowers the barrier for the ligand to diffuse into the solvent, thereby accelerating the sampling of unbinding events ^73,74^. Free energy of binding/unbinding is recovered by second-order cumulant expansion of the bias along the CVs. However, the applied bias perturbs the true dynamics, and therefore, rigorous reweighting assumptions are needed to recover the true kinetics, which may fail if the harmonic boost does not follow a Gaussian distribution or the bias becomes too large.

### 1.2 Current challenges and motivation for seekrflow

Several enhanced sampling approaches exist to improve predictions of residence times across diverse targets (Section 1.1). Supplementary Table S1 displays the performance of different enhanced-sampling approaches for drug–target residence-time predictions compared against experimental benchmarks. Although not exhaustive, it highlights successes and method-specific discrepancies, i.e., each approach excels under certain conditions. This variability motivates the development of a fully automated, end-to-end pipeline capable of estimating drug-target residence times rapidly, reproducibly, and spanning timescales from nanoseconds to hours. To summarize, estimating the kinetics and thermodynamics of drug-target binding and unbinding remains a long standing challenge due to issues including, but not limited to: (i) inadequate conformational sampling, (ii) limited simulation timescales relative to experimentally observed association and dissociation events, (iii) inaccurate force fields lacking polarization and solvent effects, (iv) integration of kinetic rate predictions with free-energy estimates within a single framework, and (v) automation of system setup, enhanced/unbiased sampling, post-simulation analysis, and error estimation. Drug-target (un)binding events comprise rare transitions, the formation of metastable states, and off-target interactions, thereby needing extensive sampling of such processes. While unbiased MD simulations require extremely long trajectories (microseconds to seconds timescales) to observe such events, enhanced sampling approaches (RaMD, GaMD, WE, and metadynamics) accelerate rare events but face issues such as distorted dynamics due to artificial biasing, limited transferability across diverse targets, choice of CVs, manual parameter tuning of biases, and recovering unbiased or true kinetics. A closely related issue is the timescale of such unbinding events across diverse targets, which often spans from a few microseconds to hours. Even with modern hardware and enhanced sampling approaches, recovering true kinetics for targets with long drug-target residence times is still impractical. Protein dynamics observed from atomistic simulations are only as accurate as the underlying force field. Failure to account for polarization at the drug-target bound state, incorrect torsional parameters, and non-bonded interactions often distort energy landscapes, leading to inaccurate kinetics. Traditional force fields rely on rigid atom typing and empirical parameters that limit their ability to represent chemical environments. To address such limitations, recent advances in machine-learned force fields, such as espaloma, which employs graph neural networks to model bonded parameters from chemical graphs, offer a differentiable and extensible alternative with improved generalizability across ligands and receptors ^75,76^. Simultaneous estimation of free energy and kinetic estimates within a single simulation framework is another major challenge. These quantities depend on distinct aspects of the free energy landscape. Thermodynamic estimates depend on state populations, while kinetic rates rely on the transition paths and barrier heights. Most methods either distort dynamics to access rare events or lack sufficient sampling to converge kinetic and thermodynamic estimates simultaneously. When drug discovery pipelines solely depend on free energy estimates, ligands with similar ΔG may have different residence times and, therefore, different drug efficacies. Computational simulation pipelines should therefore be tuned to predict thermodynamic and kinetic estimates simultaneously. The complex and multi-step process of ligand unbinding simulations lacks automation, and manual interventions are required for system setup, force field assignment, initial sampling, choice of CVs, full-length simulation, and post-simulation analyses. Such interventions limit the transferability and scalability of these methods for accelerated drug discovery campaigns. Milestoning simulations address these issues by initially employing enhanced sampling schemes to rapidly and roughly estimate the free energy landscapes, followed by the construction of milestones where independent and parallel simulations run to convergence ^45,47^. Statistical reconstruction of kinetic rates and free energy landscapes is performed via transition probability matrices derived from the flux of trajectories between milestones.

The current work presents seekrflow, which is an automated multiscale milestoning simulation pipeline that integrates unbiased atomistic simulations with machine-learned (ML) force fields. Unlike traditional approaches that rely on extensive manual setup and multiple software ecosystems, the pipeline requires only a receptor-ligand complex structure as an input, and then automates the entire workflow, i.e., from force field parameterization to running simulations to kinetic and thermodynamic analysis, providing accurate predictions of residence times and binding free energies. seekrflow is designed to reduce computational overhead while ensuring high accuracy and scalability, making it suitable for high-throughput drug discovery campaigns. This work builds upon and further extends the SEEKR ^77–79^ (Simulation-enabled estimation of kinetic rates) framework by incorporating automated milestone selection, initial choice of enhanced sampling methods, and graph neural network-based force fields for faster and accurate kinetic and thermodynamic predictions across diverse targets.

## 2 Methods

This Methods section is organized into three parts. Section 2.1 outlines the foundational principles and recent developments in the SEEKR (Simulation-enabled Estimation of Kinetic Rates) framework. Section 2.2 describes the machine-learned force field based on graph neural networks. Section 2.3 presents the design logic and implementation of seekrflow, with emphasis on accuracy, reproducibility, and scalability for high-throughput drug discovery.

### 2.1 SEEKR multiscale milestoning framework

SEEKR (Simulation-enabled Estimation of Kinetic Rates) employs a multiscale Markovian milestoning with Voronoi tessellations (MMVT) approach to partition the configurational space into Voronoi cells, with edges of these cells defined as milestones. This approach significantly reduces the computational cost of estimating drug-target kinetic rates while maintaining predictive accuracy ^79–81^. The framework estimates ligand binding and unbinding rates by independently running Brownian dynamics (BD) simulations for long-range receptor-ligand diffusion and MD simulations within Voronoi cells near the binding site, where atomistic details are essential ^79–82^.

Atomistic simulations with explicit solvation better capture interactions that govern ligand unbinding events such as hydrogen bonding, van der Waals forces, water-mediated contacts, and induced conformational changes ^83–85^. In the milestoning framework, the receptor-ligand phase space is partitioned into discrete milestones by employing Voronoi tessellations ^80,82^. Unbiased MD simulations are executed in parallel within each Voronoi cell. Milestoning theory (Supplementary Table S2) is then employed to estimate *k*_off_ by recording the statistical flux of ligand transitions between milestones ^45,77,80,86^. The last milestone, farthest from the binding site, is the kinetic boundary condition (sink state). This state defines the transition to the diffusive space where unbinding is assumed complete. In parallel, BD simulations model association kinetics (*k*_on_) employing the Browndye2 engine ^87,88^. Unlike MD, BD approximates receptor-ligand interactions with implicit solvent models and electrostatic forces, reducing computational cost while capturing essential ligand association events ^44,89–91^. BD and MD simulations are run independently within the SEEKR framework. MD simulations are further parallelized by running independent simulations within each Voronoi cell to estimate *k*_off_. Earlier versions of SEEKR implemented the NAMD MD simulation engine ^92,93^. The current version of SEEKR extends its capabilities by supporting both NAMD and OpenMM MD engines ^77,78^. This improvement enables GPU acceleration, achieves longer simulation time steps through hydrogen mass repartitioning ^94^, and provides developer-friendly flexibility for methodological advancements compared to conventional MD-based methods ^39,95,96^. With these improvements, the milestoning framework was extended to estimate the residence times of Janus kinase (JAK) inhibitors ^97^. The JAK family of proteins is central to immune signaling, with JAK2 playing a critical role in hematopoiesis and immune regulation ^98,99^. Selective inhibition of JAK2 is challenging due to its structural similarity to other JAK isoforms, especially JAK3, which shares a highly conserved adenosine triphosphate (ATP)-binding site ^100,101^. Kinetic and thermodynamic parameters for a series of tight-binding JAK2/JAK3 inhibitors were obtained employing the MMVT-SEEKR framework, and successful rank-ordering of these inhibitors based on their residence times was performed. The study demonstrated certain N-(1H-pyrazol-3-yl)pyrimidin-2-amino derivatives were selective inhibitors for JAK2 with extended residence times. Long-scale MD simulations further validated specific binding-site interactions that contributed to JAK2-specific inhibition ^97,102^.

Classical molecular mechanics (MM) force fields often struggle to accurately represent ligand-bound state interactions, such as electronic polarization and charge redistribution ^103,104^. Quantum mechanical (QM) force field reparameterization at the ligand binding site, i.e., QMrebind ^105^, addressed this issue by refining partial charges of the bound-state ligand by employing QM calculations. Milestoning simulations with QM-corrected force fields led to more accurate *k*_off_ rate predictions for therapeutically relevant complexes such as HSP90 inhibitors ^105^.

The SEEKR framework employed steered MD (SMD) simulations to generate starting structures within each Voronoi cell. A harmonic restraint is employed within SMD to gradually pull the ligand out of the binding pocket into the solvent ^78,106^, sometimes leading the ligand to follow incorrect unbinding pathways or interior surfaces of the binding pocket. Such unphysical events lead to unrealistic potential energy surfaces for milestones ^107^. Metadynamics (metaD) was incorporated into the milestoning framework to address this limitation. metaD accelerates rare event sampling by biasing the system away from previously explored states ^41^. This approach led to a more comprehensive representation of possible transition states and starting structures within Voronoi cells ^108,109^. A key application of metaD integration within SEEKR was investigating threonine-tyrosine kinase (TTK) inhibitors, an oncogenic target with prolonged inhibitor residence times ^107^. Integrating milestoning simulations with well-tempered metaD improved the ranking of inhibitors based on residence time and revealed slow conformational rearrangements affecting unbinding kinetics. metaD-initiated and milestoning-derived free energy landscapes displayed physically meaningful unbinding pathways and improved kinetic predictions.

### 2.2 Machine-Learned force field

Predicting drug-target kinetics in the milestoning framework requires precise modeling of the potential energy surfaces of the receptor-inhibitor complexes, often determined by the MM force field. Conventional force fields depend on rigid atom-typing and empirical parameterization, often limiting their ability to represent diverse chemical environments. A machine-learned, pre-trained force field model based on graph neural networks (GNNs), i.e., espaloma (extensible surrogate potential optimized by message-passing algorithms) ^75^, is integrated into the milestoning framework to address such limitations. This ML force field model replaces conventional rule-based atom typing with a fully differentiable, data-driven approach by representing molecular systems (ligands, proteins, and nucleic acids) as graphs, where atoms are nodes and bonds are edges. Unlike MM force fields that employ discrete atom-typing schemes, this learned model constructs continuous atomic embeddings through message-passing algorithms that encode local and non-local chemical environments. While the current version of the ML force field relies on fixed point charges and does not explicitly model polarization or charge transfer, its differentiable architecture enables future integration of charge polarization models.

The potential energy, *U*^MM^, of any system is given by equation 1.

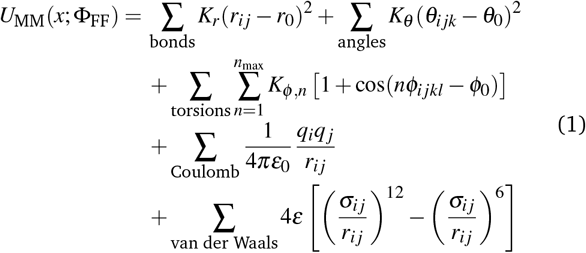

Here, the force field parameters Φ_FF_ include equilibrium bond lengths (*r*_0_), equilibrium bond angles (*θ*_0_), torsion periodicity terms (*Φ*_0_), atomic partial charges (*q*_*i*_), and Lennard-Jones parameters (*ε, σ*). To obtain these parameters, neural networks are trained on large-scale QM datasets in three stages. First, continuous atomic embeddings are generated through a message-passing neural network, where atom features are updated according to equations 2 and 3.

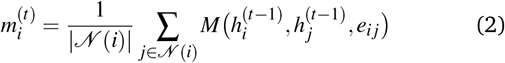

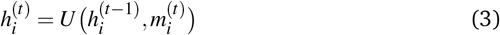

Here, 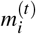 represents the aggregated messages from neighboring atoms, 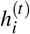 is the updated atom embedding, *N* (*i*) denotes the set of neighboring atoms bonded to atom *i*, and *M* and *U* are learnable functions. These embeddings are then employed to construct higher-order representations for molecular interactions. The rotational and permutational invariance and equivariance is achieved through symmetry-preserving pooling functions (equations 4, 5 and 6).

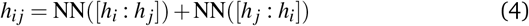

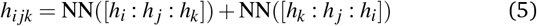

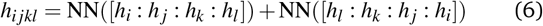

Here, *h*_*i j*_, *h*_*ijk*_, and *h*_*ijkl*_ represent bond, angle, and torsion embeddings for bonds, angles, and torsions, respectively. These embeddings are inputs to neural network functions, NN(*·*), which process atomic interactions and ensure physical consistency. The interaction terms are symmetrized to ensure invariance under atomic permutation and rotation. The force field parameters are obtained by mapping the constructed embeddings to MM parameters through fully connected neural networks. This model is trained using supervised learning with regression-based optimization that minimizes a multi-component loss function, ℒ(ΦNN) to ensure accurate reproduction of QM properties. The primary objective is to reduce the deviation between predicted and reference QM energies, as defined in equation 7.

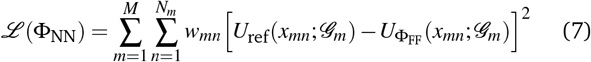

Here, *U*_ref_ represents the reference QM energy, while *U*Φ_FF_ denotes the predicted MM energy. The weighting factor *w*_*mn*_ ensures proper scaling across molecular species and conformations, preventing bias toward specific molecular classes. The loss function consists of multiple terms, incorporating energy matching, force consistency, and regularization for torsions and improper dihedral angles (equation 8).

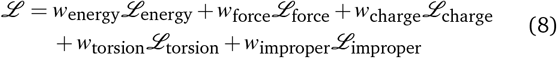

Overall, this approach results in an end-to-end differentiable force field that maintains consistency between bonded and non-bonded interactions. This force field has been validated against high-quality datasets, including drug-like small molecules, dipeptides, tripeptides, cyclic and bioactive peptides, disulfide-bridged peptides, RNA nucleosides, and trinucleotides.

### 2.3 seekrflow: Automated workflow for milestoning simulations

The current work presents seekrflow, an automated workflow for multiscale milestoning simulations to predict receptor-ligand (un)binding kinetics and thermodynamics with minimal manual intervention (Flowchart 1). Given a drug-target complex, the first step (Step 1, Flowchart 1) extracts the receptor and ligand structures into individual PDB files. This is followed by separate force field parameterization for the receptor and ligand with the espaloma pre-trained force field parameters ^75^, and the topologies of the ligand and the protein are then merged. The drug-target complex is then subjected to explicit solvation with physiological ionic strength. By default, seekrflow employs the TIP3P water model ^110^, with a solvent padding of 0.9 nm around the solute and an ionic strength of 0.15 M Na^+^*/*Cl^−^. However, the user can adjust these parameters in the seekrflow parameter file. The parameterized complex then undergoes energy minimization to resolve steric clashes. The complex topology is then saved in a PDB format, and the force field parameters are stored in XML (eXtensible Markup Language) format for subsequent MD simulations in the OpenMM MD engine ^111^.

In Step 2 (Flowchart 1), an input file with milestoning parameters is generated to automate the setup of multiscale milestoning simulations, including milestone definitions and simulation parameters for MD and BD simulations. The CV is defined as the center-of-mass (COM) distance between the ligand and alpha carbon (C*α*) atoms of the receptor within a predefined threshold (only receptor residues with C*α* atoms within this threshold distance from the ligand are included in the binding site definition with a default value of 0.8 nm). The COM-COM distance defines the anchor points along a one-dimensional reaction co-ordinate for the unbinding pathway. These anchor points then determine a set of milestone radii, partitioning the configurational space into spherical regions according to a 1D Voronoi tessellation scheme ^112^, with each milestone serving as a boundary between neighboring configurational states. An input XML file with system properties, milestone positions, and simulation parameters is then generated, which specifies the necessary information to set up system directories, configure MD and BD engines, and guide simulation steps within the milestoning framework. By default, 200 ns of MD simulations is scheduled to run for the receptor–ligand complex within each Voronoi cell. Simulations are performed using a timestep of 2 fs in the NVT ensemble at 300 K, employing Langevin dynamics with a non-bonded cutoff of 1.0 nm. For BD simulations, 1000000 trajectories are scheduled to initiate from the outermost milestone boundary (i.e., the BD region), with a grid spacing of 0.5 Å to compute the electrostatic potentials required for long-range diffusion calculations.

Step 3 (Flowchart 1) initializes the milestoning simulations by placing a receptor-ligand conformation within each Voronoi cell. Biased enhanced sampling approaches systematically explore unbinding pathways as the ligand gradually pulls out of the binding pocket. Currently, three biased sampling approaches are implemented in seekrflow. SMD ^106^ applies a harmonic restraint to gradually pull the ligand from the binding site at a predefined velocity, generating a controlled dissociation trajectory. RAMD ^113^ introduces stochastic forces to displace the ligand to explore multiple unbinding pathways. metaD ^41^ applies a history-dependent biasing potential to accelerate transitions between configurational states and improve sampling efficiency with multiple metastable states. The user can choose the preferred initial sampling approach preceding the milestoning simulations. Structures sampled from biased simulations are saved for each Voronoi cell. By default, SMD is employed with a pulling velocity of 0.01 nm ns^−1^ and a force constant of 1000 kJmol^−1^ nm^−2^ to generate dissociation trajectories along the milestones.

In Step 4 (Flowchart 1), simulation files are generated for running MD simulations in OpenMM and BD simulations in Browndye2 engines, respectively ^87,111^. System topology and force field parameters are validated for MD simulations wthin each Voronoi cell, and simulation directories are initialized to facilitate running MD simulations for each anchor. Receptor and ligand PDB files are converted to PQR molecular structure format for BD simulations ^114^. This is a requirement for the Browndye2 engine to calculate electrostatics for long-range diffusion-based association kinetics ^87^. Independent MD trajectories are run in parallel within each Voronoi cell. Reflective boundary conditions constrain the trajectories to remain within the Voronoi cells and reverse their velocities upon touching the adjacent milestone. In parallel, BD simulations are run via the Browndye2 engine to capture the long-range diffusion behavior of the ligand as it approaches the receptor.

Upon completion of milestoning and BD simulations, the final step (Step 5, Flowchart 1) aggregates the transition statistics, i.e., the number of transitions between milestones (*N*_*i j*_) and the cumulative residence times (*R*_*i*_) from all individual Voronoi cells (Supplementary Table S2). Milestoning theory constructs the transition rate matrix **Q**, and estimates the mean first passage times (MFPTs), association rate constant (*k*_on_), dissociation rate constant (*k*_off_), and free energy per milestone (Δ*G*_*i*_) (Supplementary Table S2).

**Flowchart 1.**
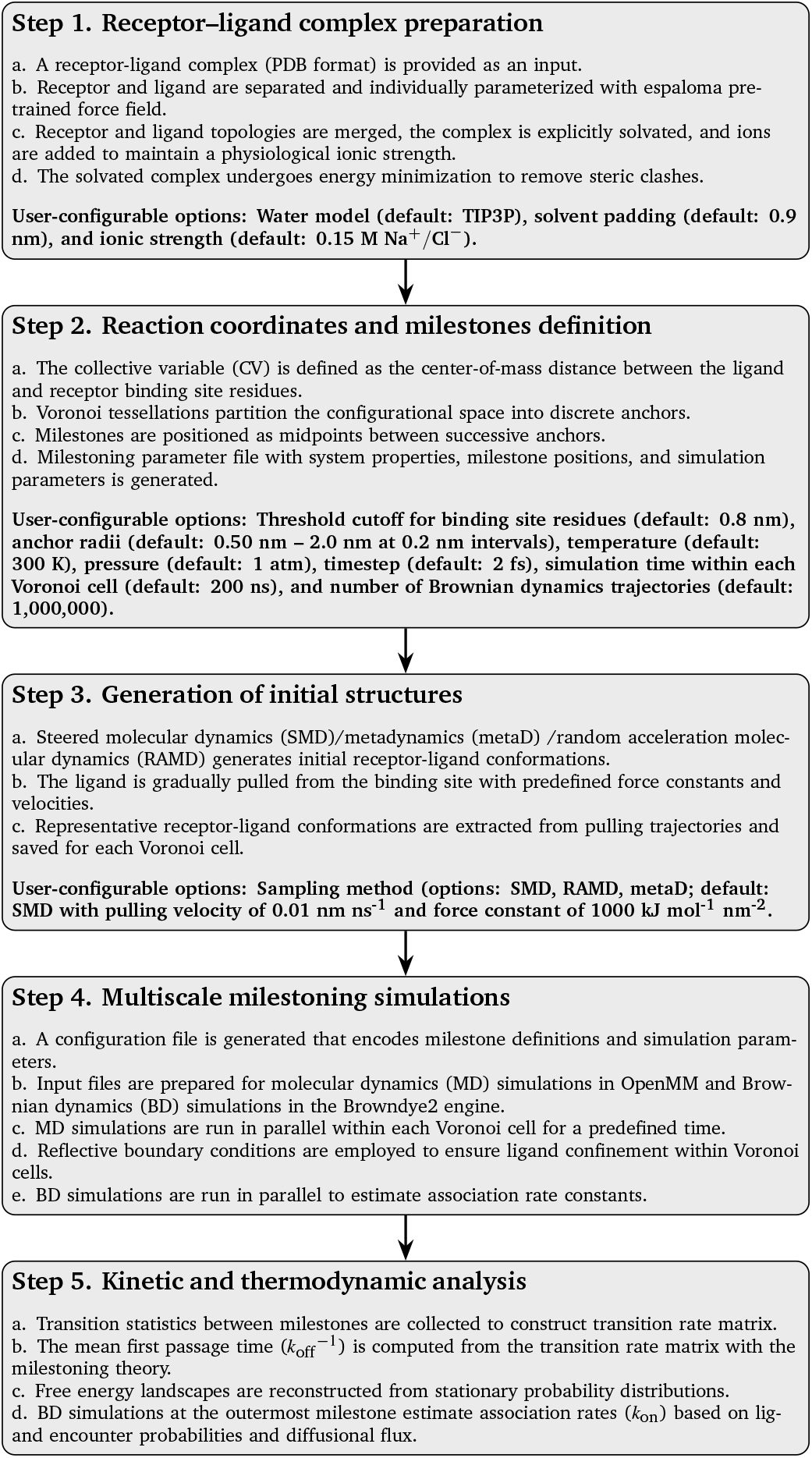
Automated milestoning simulation protocol for seekrflow.

## 3 Results

### 3.1 Trypsin-benzamidine complex

Trypsin is a serine protease that hydrolyzes peptide bonds at the carboxyl side of lysine and arginine residues, assisting in protein digestion ^115,116^ (Figure 1a). It is synthesized as an inactive trypsinogen in the pancreas but is activated in the small intestine to facilitate proteolysis and activate other digestive enzymes ^117^. Excessive trypsin activity leads to uncontrollable proteolysis, leading to pancreatic inflammation. This makes trypsin an important therapeutic target for designing inhibitors that prevent protease-driven diseases ^118,119^. Benzamidine is a reversible and competitive inhibitor of trypsin that binds to the active site through strong electrostatic and hydrogen-bonding interactions ^120,121^. Its well-defined (un)binding mode makes trypsin-benzamidine a benchmark complex to evaluate computational docking, free-energy perturbation (FEP), and enhanced simulation methods ^122–125^.

**Fig. 1.**
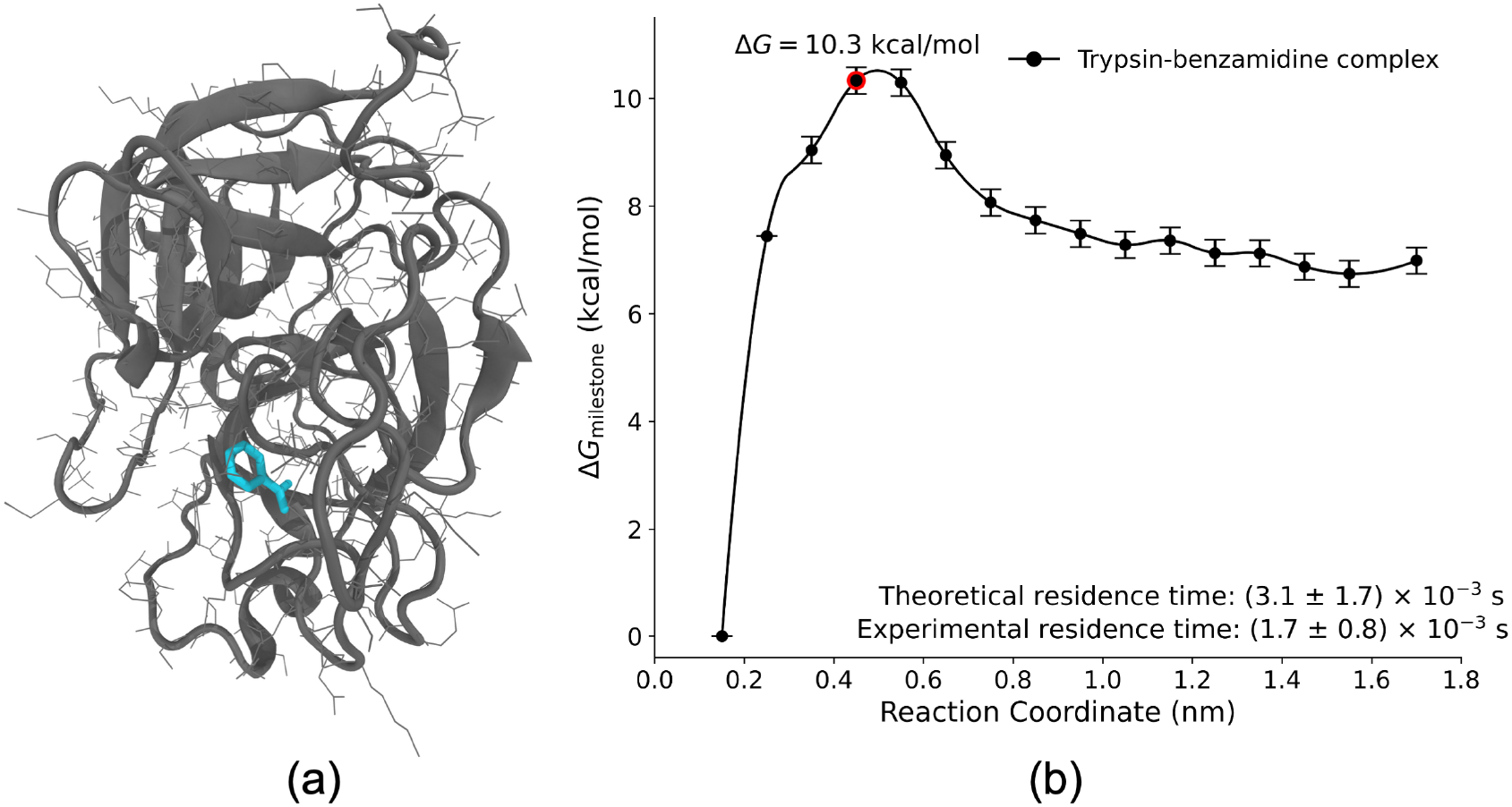
(a) Structural representation of trypsin with its inhibitor, benzamidine. (b) Free energy per milestone along the reaction coordinate for the trypsin-benzamidine complex.

Initial structure for the trypsin-benzamidine complex was obtained from our previous studies ^78,80^. The receptor and ligand were parameterized with the espaloma pre-trained force field, followed by solvation with the TIP3P water model. Na^+^*/*Cl^−^ ions were then added to maintain a physiological ionic strength of 0.15 M. The reaction coordinate for ligand unbinding was defined as the COM-COM distance between benzamidine and trypsin-binding site residues. Milestones were placed along this CV from 0.15 nm to 1.7 nm (Supplementary Table S3). The predicted residence time of (3.1 ± 1.7) × 10^−3^ s is close to the experimental residence time of (1.67 ± 0.83) × 10^−3^ s, demonstrating the accuracy of the workflow in capturing the ligand dissociation dynamics (Supplementary Table S4 and S5) ^126^. The free energy profile shows a barrier near 0.55 nm (Figure 1b), reaching a peak of approximately 10.3 kcal mol^−1^ before gradually decreasing along the unbinding pathway. Beyond this point, the free energy landscape indicates an ever-downhill dissociation and ligand unbinding is more readily achieved past this transition state.

### 3.2 HSP90-inhibitor complexes

Heat shock protein 90 (HSP90) belongs to the family of ATP-dependent molecular chaperones that are responsible for regulating the stability, activation, and maturation of many client proteins, including kinases, transcription factors, and signal molecules essential for cellular homeostasis and oncogenic transformation ^127,128^. HSP90 is a highly conserved protein across species and is expressed in various isoforms. For example, cytosolic HSP90*α* and HSP90*β*, mitochondrial TRAP1, and endoplasmic reticulum-resident GRP94 are HSP90 isoforms, each playing a distinct role in diverse cellular processes ^129,130^. HSP90 is exploited by malignant cells to support the stability and activity of mutated, overexpressed, or oncogenic proteins, thereby promoting cancer cell survival, proliferation, and metastasis ^131,132^. Unlike its latent form in normal cells, HSP90 in cancer cells exists in an activated multichaperone complex with increased ATPase activity, which makes it a critical factor in tumor progression ^127^. Its role in buffering proteotoxic stress also enables cancer cells to tolerate oncogenic mutations, making it a prime therapeutic target ^133^. HSP90 inhibitors function by blocking ATP binding and hydrolysis at the N-terminal domain, preventing the formation of functional HSP90-client complexes, leading to the degradation of oncogenic client proteins via the ubiquitin-proteasome system, ultimately inducing apoptosis in cancer cells (Supplementary Figure S1) ^134^.

Given its involvement in multiple hallmarks of cancer, including sustained proliferative signaling, evasion of apoptosis, and enhanced angiogenesis, HSP90 has emerged as an attractive target for cancer therapy ^135,136^. Highly sensitive HSP90 clients such as human epidermal growth factor receptor 2 (HER2), epidermal growth factor receptor (EGFR), and anaplastic lymphoma kinase (ALK), have shown promising responses to HSP90 inhibition ^137–140^. Despite significant progress, challenges remain in optimizing HSP90 inhibitors for clinical use. Off-target toxicity, resistance mechanisms, and the compensatory heat shock response, which upregulates cytoprotective proteins such as HSP70, limit the long-term efficacy of HSP90 inhibitors ^141,142^.

The N-terminal domain of HSP90 has a highly flexible ATP-binding site that undergoes loop/helix rearrangements upon ligand binding ^143^. Enhanced sampling simulations have struggled to capture the kinetics of such slow conformational changes, as different inhibitors can induce local rearrangements of the ATP binding site (loop-in, loop-out, or helical states). Moreover, due to the existence of multiple ligand unbinding pathways, hidden allosteric couplings, and unbinding kinetics spanning from seconds to hours, there exist severe sampling bottlenecks to study HSP90-inhibitor complexes ^144,145^.

HSP90 inhibitors in this study were chosen based on their scaffold diversity, range of experimentally-determined residence times, and binding preference for either loop or helix conformations (Figure 2a). The subset of inhibitors includes resorcinol, hydroxy-indazole, benzamide, aminoquinazoline, aminopyrrolopyrimidine, and 2-aminopyramidine scaffolds (Supplementary Table S6). The binding mode of HSP90 inhibitors varies based on their interaction with the N-terminal ATP-binding pocket ^146,147^. Some inhibitors bind within the ATP site and stabilize the flexible loop region that undergoes conformational rearrangement upon ligand binding (loop conformation) ^148^. Other inhibitors interact with the transient hydrophobic subpocket near *α*-helix3 beyond the ATP pocket, which leads to additional stabilization of the complex (helix conformation) ^36,149,150^. HSP90 inhibitors in this study span residence times from a few seconds to hours (Supplementary Tables S7 and S8), which allows us to evaluate the reliability of the automated milestoning pipeline in predicting distinct kinetic profiles.

**Fig. 2.**
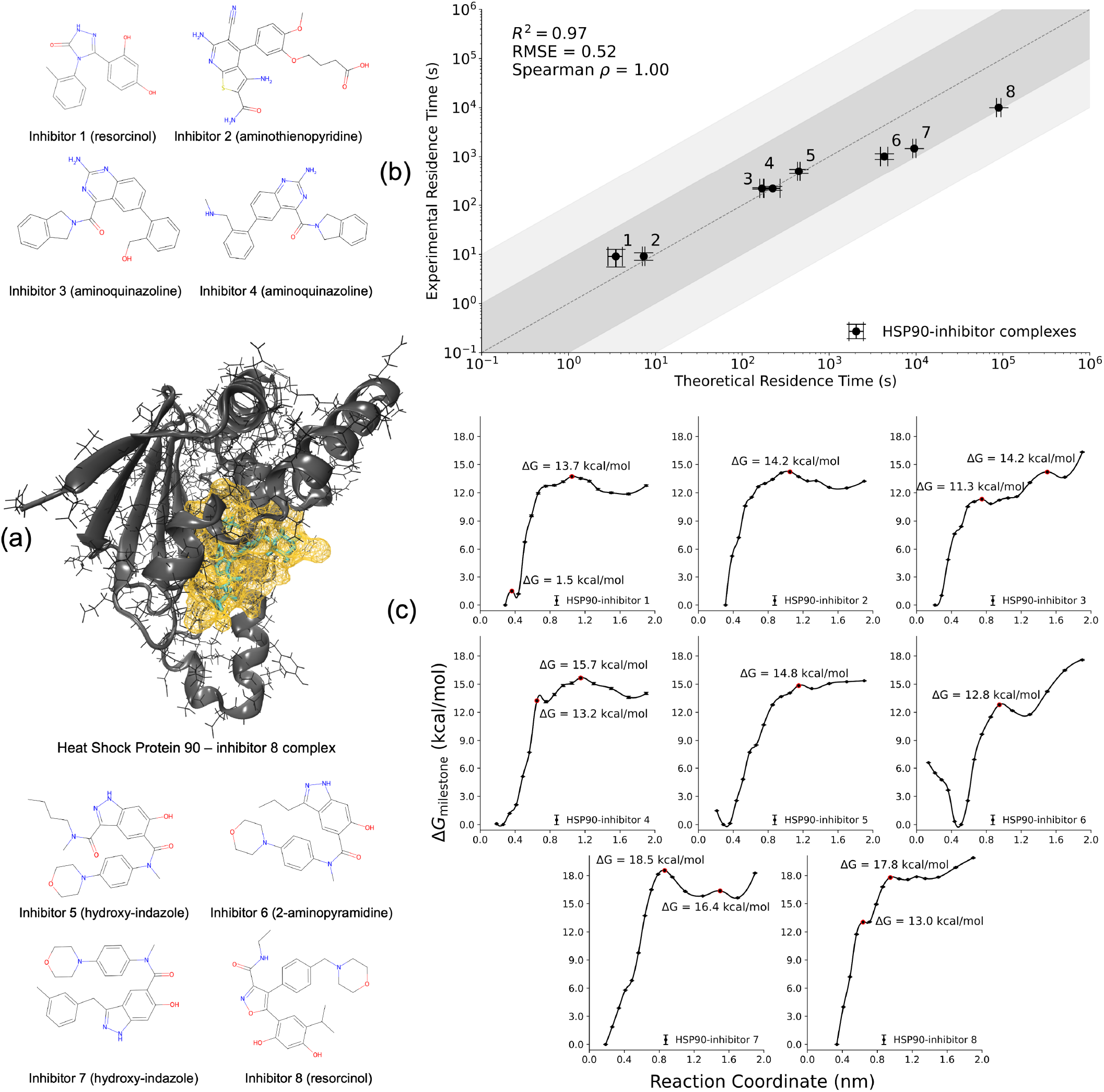
(a) Structural representation of heat shock protein 90 (HSP90) and its inhibitors. The chemical structures of eight HSP90 inhibitors, including resorcinol, aminothienopyridine, aminoquinazoline, hydroxy-indazole, and 2-aminopyramidine derivatives, are represented. (b) Experimental and predicted residence times for HSP90-inhibitor complexes. The shaded regions indicate deviations within one (dark gray) and two (light gray) orders of magnitude. Regression analysis performed in the log_10_ space yields *R*^2^ = 0.97, RMSE = 0.52, and Spearman correlation coefficient = 1.0. (c) Free energy profiles of HSP90-inhibitor complexes from multiscale milestoning simulations. The x-axis represents the center-of-mass (COM) distance between the ligand and binding site, and the y-axis shows free energy in kcal mol−1. Transition-state barriers and inflection points (intermediate metastable states) are indicated by red dots. Error bars denote statistical uncertainties 78.

Initial structures for the HSP90-inhibitor complexes were obtained from our previous study ^36,105,151–153^. The current workflow employs a standardized set of scripts common to all complexes and executes them sequentially with minimal manual intervention to obtain kinetic and thermodynamic parameters. For each complex, the receptor and ligand were separated into individual PDB files and parameterized with the pre-trained espaloma force field model. Complexes were solvated with the TIP3P water model, and Na^+^*/*Cl^−^ ions were added at a concentration of 0.15 M to mimic physiological ionic strength. The CV was defined as the distance between the COM of ligand atoms and the C*α* atoms of the binding site receptor residues, with a predefined threshold of 8 Å (only receptor residues with C*α* atoms within this distance from the ligand were included in the binding site definition). Anchors were placed at increasing distances along the CV, and milestones were placed at midpoints between the successive anchors (Supplementary Table S9).

To ensure comprehensive initial sampling within each Voronoi cell, the holo insertion by directed restraints (HIDR) method was adopted to generate initial structures. HIDR applies a restraint force constant and a default translation velocity of 0.05 nm ns^−1^ for SMD simulations to gradually pull the ligand towards each target anchor while avoiding non-physical paths. SMD-generated conformations were then adopted as starting structures for parallel milestoning simulations. MD simulations were run for 200 ns within each Voronoi cell for 16 cells, amounting to 3.2 *µ*s of simulation time per complex. All simulations were run with default parameters to ensure a consistent and automated workflow from structure preparation to kinetic/thermodynamic estimates.

The experimental and predicted *k*_off_ estimates for the HSP90-inhibitor complexes show an excellent agreement, with all estimates falling within a single order of magnitude of experimental measurements (Figure 2b). This high level of accuracy is reflected in the regression analysis performed in the log_10_ space, yielding *R*^2^ of 0.97, RMSE of 0.52, and a Spearman correlation coefficient *ρ* of 1.0. Accurate *k*_off_ predictions for inhibitors with longer experimental residence times (*k*_off_ *<* 10^−4^ *s*^−1^) using default parameters without augmentation (QMrebind for force field correction or metaD for initial sampling) demonstrates the ability of the automated milestoning approach to capture slow dissociation kinetics accurately. The rank ordering of residence times for the series of inhibitors is well preserved. Some deviations are observed for inhibitors with longer residence times (Figure 2b), where small errors in *k*_off_ estimation can lead to larger discrepancies in residence time estimates due to their inverse relationship. Despite this, the overall trend remains consistent with experimental observations, demonstrating the accuracy of the approach.

Figure 2c depicts the free energy landscapes of HSP90-inhibitor unbinding. Fast dissociating inhibitors (HSP90-inhibitors 1 and 2) exhibit relatively shallow free energy barriers along the CV, with minimal intermediate metastable states, thus implying a more direct and quicker dissociation pathway. Tight binding inhibitors (HSP90-inhibitor 8) exhibit higher free energy barriers and multiple inflection points corresponding to transient intermediates that stabilize the complex before dissociation. The higher free energy barriers correlate with large inhibitor residence times, explaining the thermodynamic origin of their kinetic stability. HSP90-inhibitors 6 and 7 bind with the helix conformation of the ATP-binding pocket, where the hydrophobic *α*-helix3 subpocket is open. Such interactions result in complex free energy profiles with high energy barriers and multiple intermediates. Additional stabilizing interactions within this subpocket potentially contribute to longer residence times compared to inhibitors that bind exclusively within the ATP pocket. Uniform milestone placements across complexes ensured that differences in the energy profiles stem from inherent receptor-ligand interactions, not milestone positioning variability. In conclusion, thermodynamic trends for HSP90-inhibitor complexes correlate with the kinetic profiles, favoring the involvement of free energy barriers and metastable states in explaining inhibitor residence times.

### 3.3 TTK-inhibitor complexes

Threonine tyrosine kinase (TTK) or monopolar spindle 1 (Mps1) is a dual specificity protein kinase that plays an important role in mitosis checkpoint signaling. TTK regulates the spindle assembly checkpoint (SAC), a surveillance mechanism that ensures proper chromosome segregation during mitosis ^154–156^. It prevents chromosome missegregation and maintains genomic stability by phosphorylating SAC proteins, facilitating the recruitment of mitotic checkpoint complex (MCC). This complex then suppresses the anaphase-promoting complex/cyclosome (APC/C) activity until chromosomes attach correctly to the spindle microtubules ^157,158^. TTK further ensures proper microtubule-kinetochore attachments through phosphorylation of key substrates. Once all kinetochores are correctly aligned, TTK is removed, SAC is deactivated, MCC disassembles, and APC/C becomes active, initiating anaphase progression ^159,160^.

Overexpression of TTK is observed in many malignant tumors, including breast cancer, glioblastoma, hepatocellular carcinoma, and pancreatic cancer ^161,162^. Since its overexpression is correlated with poor prognosis, TTK is a promising therapeutic target. Inhibition of TTK disrupts SAC function, causing premature mitotic exit, chromosome missegregation, and aneuploidy ^163,164^. Despite its therapeutic potential, TTK inhibitor faces multiple challenges. For example, inhibitor resistance from mutations in the ATP-binding pocket reduces the ligand binding affinity and limits the effectiveness of these inhibitors ^165,166^. Off-target toxicity is a major concern since TTK performs essential functions in normal cell proliferation processes, thus calling for the design of selectivity enhancement strategies. Therefore, drug residence time must be optimized to achieve maximal antiproliferative activity while minimizing adverse outcomes ^167,168^.

Computational modeling of TTK inhibitors face significant challenges due to the structural variability and flexibility of key domain residues, such as glycine-rich gating (G-loop) and activation loops. Recent studies have demonstrated the formation of multiple metastable states upon inhibitor binding ^107,169–171^. The unbinding mechanism is also influenced by large-scale protein conformational changes and solvent effects ^169^. Conventional and enhanced sampling approaches fail to achieve accurate free energy landscapes and unbinding kinetics due to insufficient exploration of high-dimensional conformational spaces such as coupled dynamics, solvent-mediated interactions, and ligand orientations. The current pipeline addresses sampling limitations by first exploring the phase space with enhanced sampling schemes such as metadynamics or steered MD, followed by unbiased MD simulations within milestones to reconstruct the unbinding kinetics.

TTK inhibitors were selected for their structural and functional diversity and known challenges in kinetic modeling due to their complex unbinding pathways involving transient binding poses and potential allosteric effects (Figure 3a) ^169^. These inhibitors demonstrated a range of experimental residence times, from seconds to hours. Structural data for TTK-inhibitor complexes were derived from a prior study that incorporated high-resolution crystal structures as the primary source ^107,170–173^. Similar protocols were followed for preparing TTK-inhibitor complexes as those for the HSP90-inhibitor complexes. The receptor and ligand were saved into individual PDB files, followed by parameterization with the espaloma pre-trained force field model and solvation with the TIP3P water model at a physiological ionic strength of 0.15 M Na^+^*/*Cl^−^. The CV, defined as the COM-COM distance between ligand atoms and the C*α* atoms of binding site residues with a threshold of 8 Å, was used to determine anchor positions (Supplementary Table S10). Initial sampling within Voronoi cells was performed using the HIDR approach that generated starting structures for milestoning simulations via SMD simulations. Parallel MD simulations were run for 200 ns within each Voronoi cell for 16 cells, totaling 3.2 *µ*s of simulation time per complex.

**Fig. 3.**
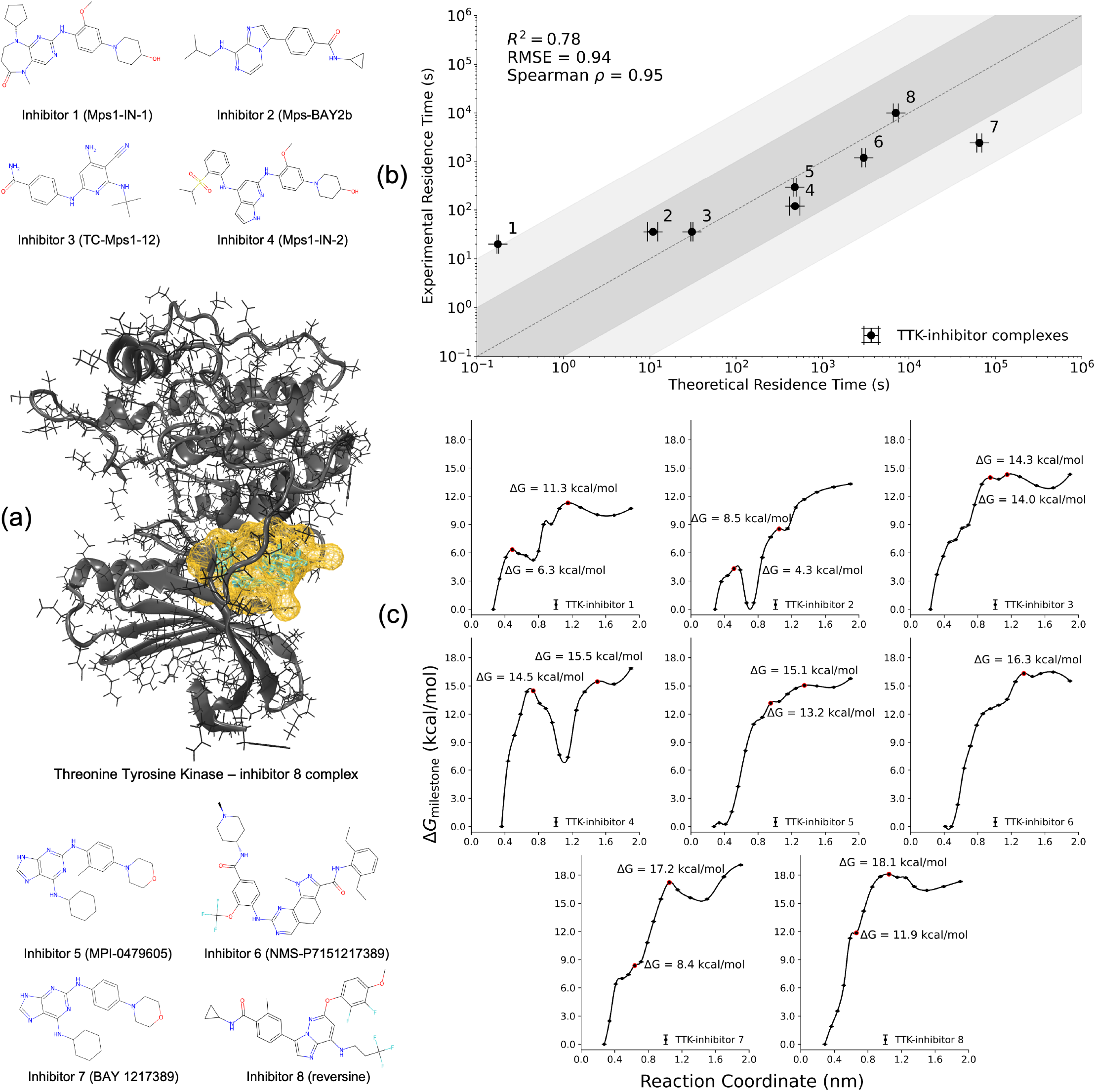
(a) Structural representation of threonine tyrosine kinase (TTK) and its inhibitors. The chemical structures of eight TTK inhibitors, including Mps1-IN-1, Mps-BAY2b, TC-Mps1-12, Mps1-IN-2, MPI-0479605, NMS-P715, BAY 1217389, and reversing, are represented. (b) Experimental and predicted residence times for TTK-inhibitor complexes. The shaded regions indicate deviations within one (dark gray) and two (light gray) orders of magnitude. Regression analysis performed in the log_10_ space yields *R*^2^ = 0.78, RMSE = 0.94, and Spearman correlation coefficient = 0.95. (c) Free energy profiles of TTK-inhibitor complexes from multiscale milestoning simulations. The x-axis represents the center-of-mass (COM) distance between the ligand and binding site, and the y-axis shows free energy in kcal mol−1. Transition-state barriers and inflection points (intermediate metastable states) are indicated by red dots. Error bars denote statistical uncertainties 78.

The experimental and predicted *k*_off_ rates for TTK-inhibitor complexes show a strong correlation, with the majority of the predictions falling within a single order of magnitude of the experimental data (Figure 3b, Supplementary Table S11 and S12). Regression analysis in the log_10_ space (*R*_2_ = 0.78, RMSE = 0.94, and Spearman correlation coefficient, *ρ* = 0.95) demonstrates the accuracy of the automated milestoning framework in estimating the kinetics and rank ordering of the TTK-inhibitor complexes. Note that all simulations were run for TTK-inhibitor complexes with default parameters. Predicted residence time estimates preserve the rank order observed in experimental estimates across inhibitors. Free energy profiles of TTK-inhibitor complexes display distinct thermodynamic features correlating with their residence times (Figure 3c). TTK-inhibitor 1, with the shortest residence time, has a low free energy barrier and a smooth unbinding pathway with minimal resistance to dissociation. For tight binding inhibitors such as TTK inhibitors 6, 7, and 8, free energy barriers become more prominent with multiple intermediates. TTK inhibitors 3 and 4, with multiple peaks in their free energy landscapes, suggest complex unbinding pathways with intermediate states that may lead to transient inhibitor stabilization before complete dissociation.

## 4 Discussion

Accurately predicting ligand (un)binding kinetics remains a central challenge in computational drug discovery. This necessitates approaches that balance efficiency, accuracy, and automation. The automated multiscale milestoning pipeline presented in this study advances the state-of-the-art simulation methods by integrating ML force fields and enhanced sampling methods within a highly parallelizable framework. Current methodologies often rely on manually intensive workflows or employ empirical force fields for complex parameterization with limited transferability. The current approach ensures automation and accuracy with the potential to facilitate high-throughput screening and lead optimization with minimal manual intervention.

Conventional MD simulations require excessively long simulation timescales ranging from milliseconds to hours to capture spontaneous unbinding events. This makes conventional atomistic simulations impractical for high-throughput screening of drug candidates. Enhanced sampling methods such as metaD, SMD, and RAMD ^35,106,109^ expedite unbinding events. Despite their computational efficiency, such methods often distort natural unbinding pathways. Reproducibility remains a concern, as the observed unbinding events may vary depending on simulation parameters, initial conditions, and the applied external bias, making it difficult to obtain consistent kinetic predictions across independent runs. These methods also require extensive prior knowledge about CVs and careful parameter tuning, such as pulling velocity in SMD or the random force magnitude in RAMD. In metaD simulations, the Gaussian deposition rate and width must be set appropriately to facilitate enhanced sampling without artificially modifying transition-state barriers ^109^. Improper selection of such parameters may lead to non-physical unbinding mechanisms, biased free energy landscapes, and inconsistent kinetic predictions.

The seekrflow framework bypasses potential issues described above by establishing an automated multiscale milestoning strategy that partitions configurational space into milestones and performs unbiased MD simulations followed by thermodynamic profiling and kinetic rate estimations for receptor-ligand complexes. To date, milestone placements, choice of CVs, force field parameterization, and atomistic simulations have been handled separately, which required a careful manual setup. The current workflow integrates all these steps into an automated workflow, i.e., from structure preparation to kinetic and thermodynamic analysis, thereby eliminating manual intervention for accelerated drug discovery campaigns. The framework accurately estimated kinetic and thermodynamic parameters across multiple receptor-ligand complexes, from trypsin-benzamidine complex to more pharmacologically relevant targets, including HSP90 and TTK proteins. Predicted residence times show a high correlation with experimental values (Figures 2, 3, and 1), with most estimates falling within one order of magnitude of the experimental estimates. The pipeline also preserves the rank-ordering of residence times, a key requirement for rational drug design. Figure 4 shows the steady progress of the SEEKR milestoning scheme from host-guest complexes to more complex targets such as HSP90 and TTK proteins, demonstrating consistent estimation accuracy across increasing drug-target complexity.

**Fig. 4.**
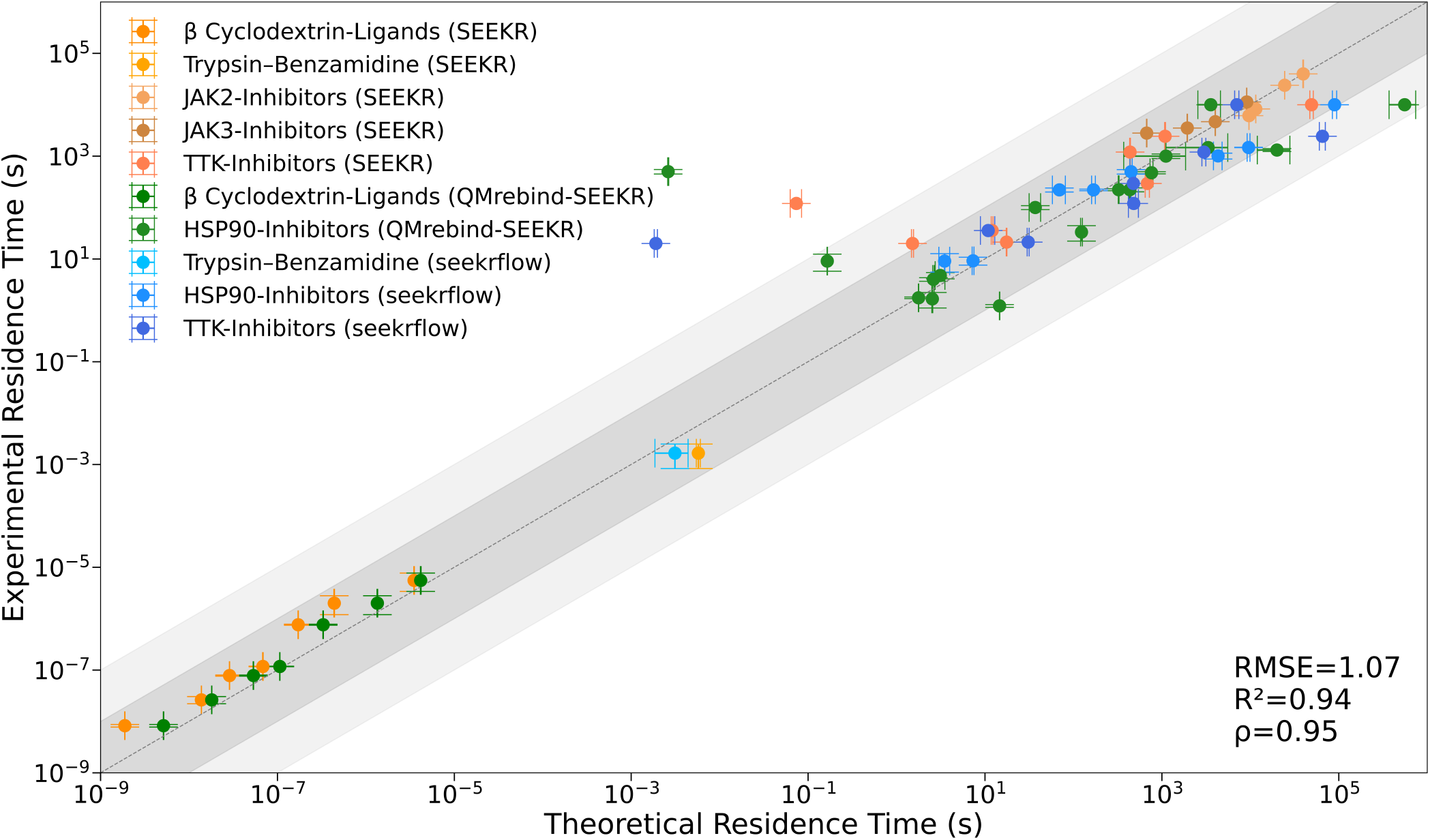
Experimental and SEEKR-predicted residence times for multiple receptor-ligand complexes using conventional force fields, including model host-guest complexes, trypsin-benzamidine complex, and therapeutically relevant targets such as HSP90, TTK, and JAK-inhibitor complexes. The dark gray shaded area represents predictions within one order of magnitude of the experimental values, while the light gray shaded area represents predictions within two orders of magnitude. Regression analysis performed in the log_10_ space yields RMSE = 1.07, R2 = 0.94, and Spearman correlation coefficient = *ρ* = 0.95.

A key advantage of seekrflow is its scalability as it achieves near-ideal parallel efficiency by partitioning the configurational space into independent milestones, reducing computational costs and simulation time. Such enhancements enable accurate kinetic predictions for ligands with residence times from milliseconds to hours, making them particularly relevant for tight-binding inhibitors. Despite its advantages, limitations remain. While ML force fields improve unbinding rate estimates, further refinements may better capture bound-state polarization effects ^105,174^. ML-driven CVs could refine milestone selection to better represent complex energy landscapes of ligand unbinding pathways. Beyond drug discovery, the methodology outlined in this work is broadly applicable to diverse biomolecular interactions, including protein-protein and protein-DNA (un)binding kinetics. The ability of this framework to model long-timescale biomolecular interactions with high accuracy holds promise for applications in systems biology, enzyme engineering, and biomaterials design. Integrating the seekrflow multiscale milestoning pipeline with experimental approaches such as cryo-electron microscopy, X-ray crystallography, and NMR spectroscopy can further enhance the understanding of bound and unbound (apo-holo) target states for the discovery of novel drugs ^175–178^. Experimental observables added as constraints or inputs to the simulation pipeline further refine the accuracy of the pipeline and establish a more robust connection between the sequence, structure, dynamics, and kinetics.

Despite its success in accurately predicting absolute kinetics (and thermodynamics) for a wide range of protein-ligand complexes, the milestoning simulation pipeline has certain limitations and trade-offs. To date, a relatively straightforward CV (COM-COM distance between the ligand and the C*α* atoms of the binding site) defines the milestone placements. This could potentially be an issue when the true unbinding pathway of the ligand involves additional variables, such as a particular rotation or a side-chain movement. Accounting for additional degrees of freedom may require multidimensional milestoning at the cost of increased computational expense. The pipeline could be extended to automatically and adaptively accommodate additional milestones. For example, if the ligand spends an unusually long time within a particular milestone with scarce transitions to its adjacent milestones, an additional milestone could adaptively be inserted at the bottleneck.

The current protocol is designed for broad applicability, i.e., for a wide range of protein-ligand complexes. This one-size-fits-all approach may sometimes need careful manual tuning for optimal milestone placement. For example, the milestones must be manually tuned with well-defined CVs in certain protein-ligand complexes, where multiple ligand-exiting pathways are observed ^179–181^. In cases where flexible proteins or allosteric binding sites are involved, modeling ligand unbinding processes may involve accounting for slow protein rearrangements ^182–184^, such as loop and domain motions, that are not fully accounted for in the current protocol. While the milestoning simulations adequately capture the motion of the ligand in the phase space, it also assumes the quasi-static environment of the protein within each milestone. Large conformational changes could violate such an assumption, and a fine-tuned CV would then be needed to capture such motions adequately. Alternatively, adaptive milestoning placements based on extensive initial sampling could account for crucial degrees of freedom for the protein conformational changes.

The accuracy of residence time predictions depends largely on the quality of the force fields. Conventional force fields may introduce artifacts. For instance, overestimating ligand interactions with the binding site can cause unbinding to occur too slowly, or missing electronic polarization at the binding site can lead to rapid unbinding. Recent studies with quantum-corrected force fields at the ligand-bound state accounted for accurate *k*_off_ predictions for HSP90 inhibitors ^105^. Future improvements in milestoning simulations may include ML-force fields incorporating ligand-binding site polarization effects informed by quantum mechanical calculations.

## 5 Conclusion

Building upon existing multiscale milestoning approaches, the current study introduces a fully automated simulation pipeline that integrates ML force fields with enhanced sampling methods. In this way, we reinforce the existing advantages of conventional milestoning simulations while introducing improvements in computational efficiency and accuracy. This framework enables unbiased atomistic simulations to effectively capture rare unbinding events and estimate kinetic and thermodynamic predictions for a range of receptor-inhibitor complexes. seekrflow automates every stage of the process, i.e., from receptor-ligand parameterization to milestone definition and kinetic/thermodynamic analysis. Incorporating ML force fields trained on QM data addresses certain limitations of classical force fields. The compatibility of ML fields with the OpenMM MD engine further simplifies the parameterization process and ensures an efficient integration into the simulation pipeline. The multiscale milestoning approach efficiently scales to account for complex systems with a wide range of residence times, executes parallel simulations, and reduces computational load compared to traditional methods. Our results demonstrate strong agreement with experimental data across diverse receptor-ligand complexes, making the pipeline an effective and reliable tool for drug discovery. This pipeline sets a new standard for physics-based, high-throughput kinetic and thermodynamic predictions and advances the design of next-generation therapeutics.

## Acknowledgement

The authors thank Nehemiah Zewde (Genentech, Inc.) for his valuable insights and discussions that contributed to this work. The authors acknowledge Javier O. Sanlley-Hernandez for his testing of the seekrflow code. The authors acknowledge Open-Eye Scientific Software for providing academic licenses to use their tools. The Flatiron Institute is a division of the Simons Foundation.

## Supplementary information

The supplementary information file includes theoretical and experimental unbinding rates, residence times, and milestone definitions for TTK-inhibitor, HSP90-inhibitor, and trypsin-benzamidine complexes.

## Data and Software Availability

The seekrflow automated milestoning pipeline for computing receptor-ligand kinetics and thermodynamics is available at https://github.com/seekrcentral/seekrflow, and the SEEKR2 software can be accessed at https://github.com/seekrcentral/seekr2. All data supporting this study, including milestoning simulations for eight HSP90-inhibitor complexes, eight TTK-inhibitor complexes, and the trypsin-benzamidine complex, along with structural and topology files, simulation trajectories, and analysis scripts for computing receptor-ligand unbinding rates, are available as a supplemental dataset at https://zenodo.org/uploads/14968047.

## References

1 A. S. Powers, V. Pham, W. A. Burger, G. Thompson, Y. Laloudakis, N. W. Barnes, P. M. Sexton, S. M. Paul, A. Christopoulos, D. M. Thal et al., Nature Chemical Biology, 2023, 19, 805–814.

2 I. Guryanov, S. Fiorucci and T. Tennikova, Materials Science and Engineering: C, 2016, 68, 890–903.

3 W. J. van der Velden, L. H. Heitman and M. M. Rosenkilde, ACS pharmacology & translational science, 2020, 3, 179–189.

4 D. A. Schuetz, W. E. A. de Witte, Y. C. Wong, B. Knasmueller, L. Richter, D. B. Kokh, S. K. Sadiq, R. Bosma, I. Nederpelt, L. H. Heitman et al., Drug discovery today, 2017, 22, 896–911.

5 K. E. Knockenhauer and R. A. Copeland, British Journal of Pharmacology, 2024, 181, 4103–4116.

6 M. Srinivasarao and P. S. Low, Chemical reviews, 2017, 117, 12133–12164.

7 G. Vauquelin and S. J. Charlton, British journal of pharmacology, 2010, 161, 488–508.

8 J. Pettinger, M. Carter, K. Jones and M. D. Cheeseman, Journal of Medicinal Chemistry, 2019, 62, 11383–11398.

9 B. Han, F. G. Salituro and M.-J. Blanco, ACS Medicinal Chemistry Letters, 2020, 11, 1810–

10 X. Xie, T. Yu, X. Li, N. Zhang, L. J. Foster, C. Peng, W. Huang and G. He, Signal transduction and targeted therapy, 2023, 8, 335.

11 R. Abel, L. Wang, E. D. Harder, B. Berne and R. A. Friesner, Accounts of chemical research, 2017, 50, 1625–1632.

12 Z. Cournia, B. Allen and W. Sherman, Journal of chemical information and modeling, 2017, 57, 2911–2937.

13 E. King, E. Aitchison, H. Li and R. Luo, Frontiers in molecular biosciences, 2021, 8, 712085.

14 M. Bernetti, A. Cavalli and L. Mollica, MedChemComm, 2017, 8, 534–550.

15 S. R. Hoare, B. A. Fleck, J. P. Williams and D. E. Grigoriadis, Drug Discovery Today, 2020, 25, 7–14.

16 H. Lu and P. J. Tonge, Current opinion in chemical biology, 2010, 14, 467–474.

17 M. R. Dowling and S. J. Charlton, British journal of pharmacology, 2006, 148, 927–937.

18 G. K. Walkup, Z. You, P. L. Ross, E. K. Allen, F. Daryaee, M. R. Hale, J. O’donnell, D. E. Ehmann, V. J. Schuck, E. T. Buurman et al., Nature chemical biology, 2015, 11, 416–423.

19 P. Carpinelli, R. Ceruti, M. L. Giorgini, P. Cappella, L. Gianellini, V. Croci, A. Degrassi, G. Texido, M. Rocchetti, P. Vianello et al., Molecular cancer therapeutics, 2007, 6, 3158– 3168.

20 R. Zhang and W. T. Windsor, Antiviral methods and protocols, 2013, 59–79.

21 D. Kitagawa, M. Gouda and Y. Kirii, Journal of Biomolecular Screening, 2014, 19, 453–461.

22 C. S. Tautermann, T. Kiechle, D. Seeliger, S. Diehl, E. Wex, R. Banholzer, F. Gantner, M. P. Pieper and P. Casarosa, Journal of medicinal chemistry, 2013, 56, 8746–8756.

23 R. A. Copeland, Nature Reviews Drug Discovery, 2016, 15, 87–95.

24 L. Wang, J. Chambers and R. Abel, Biomolecular simulations: methods and protocols, Springer, 2019, pp. 201–232.

25 L. F. Song, T.-S. Lee, C. Zhu, D. M. York and K. M. Merz Jr, Journal of chemical information and modeling, 2019, 59, 3128–3135.

26 S. Genheden and U. Ryde, Expert opinion on drug discovery, 2015, 10, 449–461.

27 T. J. Lane, D. Shukla, K. A. Beauchamp and V. S. Pande, Current opinion in structural biology, 2013, 23, 58–65.

28 C. Schütte, S. Klus and C. Hartmann, Acta Numerica, 2023, 32, 517–673.

29 G. Henkelman, H. Jónsson, T. Lelièvre, N. Mousseau and A. F. Voter, Handbook of Materials Modeling: Methods: Theory and Modeling, 2020, 825–834.

30 P. Conflitti, S. Raniolo and V. Limongelli, Journal of chemical theory and computation, 2023, 19, 6047–6061.

31 Y. Miao, C.-E. A. Chang, W. Zhu and J. A. McCammon, Mechanisms, thermodynamics and kinetics of ligand binding revealed from molecular simulations and machine learning, 2023.

32 J. Wang, Q. Wang and S. Song, Journal of Chinese Pharmaceutical Sciences, 2020, 29, year.

33 G. Spoto and M. Minunni, The journal of physical chemistry letters, 2012, 3, 2682–2691.

34 R. Zhang, C. M. Barbieri, M. Garcia-Calvo, R. W. Myers, D. McLaren and M. Kavana, Front Biosci (Schol Ed), 2016, 8, 278–297.

35 S. Kalyaanamoorthy and Y.-P. P. Chen, Journal of chemical information and modeling, 2012, 52, 589–603.

36 D. B. Kokh, M. Amaral, J. Bomke, U. Gradler, D. Musil, H.-P. Buchstaller, M. K. Dreyer, M. Frech, M. Lowinski, F. Vallee et al., Journal of chemical theory and computation, 2018, 14, 3859–3869.

37 Y. Miao, The Journal of chemical physics, 2018, 149, year.

38 L. Mollica, S. Decherchi, S. R. Zia, R. Gaspari, A. Cavalli and W. Rocchia, Scientific reports, 2015, 5, 11539.

39 L. Mollica, I. Theret, M. Antoine, F. Perron-Sierra, Y. Charton, J.-M. Fourquez, M. Wierzbicki, J. A. Boutin, G. Ferry, S. Decherchi et al., Journal of medicinal chemistry, 2016, 59, 7167–7176.

40 W. Sinko, Y. Miao, C. A. F. de Oliveira and J. A. McCammon, The Journal of Physical Chemistry B, 2013, 117, 12759–12768.

41 A. Barducci, M. Bonomi and M. Parrinello, Wiley Interdisciplinary Reviews: Computational Molecular Science, 2011, 1, 826–843.

42 D. M. Zuckerman and L. T. Chong, Annual review of biophysics, 2017, 46, 43–57.

43 S.-H. Ahn, A. A. Ojha, R. E. Amaro and J. A. McCammon, Journal of chemical theory and computation, 2021, 17, 7938–7951.

44 G. A. Huber and S. Kim, Biophysical journal, 1996, 70, 97–110.

45 A. A. Ojha, L. W. Votapka and R. E. Amaro, Journal of Chemical Theory and Computation, 2024, 20, 9759–9769.

46 R. Elber, A. Fathizadeh, P. Ma and H. Wang, Wiley Interdisciplinary Reviews: Computational Molecular Science, 2021, 11, e1512.

47 R. Elber, Annual review of biophysics, 2020, 49, 69–85.

48 B. E. Husic and V. S. Pande, Journal of the American Chemical Society, 2018, 140, 2386– 2396.

49 J. D. Chodera and F. Noé, Current opinion in structural biology, 2014, 25, 135–144.

50 A. A. Ojha, S. Thakur, S.-H. Ahn and R. E. Amaro, Journal of chemical theory and computation, 2023, 19, 1342–1359.

51 K. Lindorff-Larsen, S. Piana, R. O. Dror and D. E. Shaw, Science, 2011, 334, 517–520.

52 J. L. Klepeis, K. Lindorff-Larsen, R. O. Dror and D. E. Shaw, Current opinion in structural biology, 2009, 19, 120–127.

53 Y. Shan, E. T. Kim, M. P. Eastwood, R. O. Dror, M. A. Seeliger and D. E. Shaw, Journal of the American Chemical Society, 2011, 133, 9181–9183.

54 N. Plattner and F. Noé, Nature communications, 2015, 6, 7653.

55 J. Wang, H. N. Do, K. Koirala and Y. Miao, Journal of Chemical Theory and Computation, 2023, 19, 2135–2148.

56 M. Sarich, F. Noé and C. Schütte, Multiscale Modeling & Simulation, 2010, 8, 1154–1177.

57 R. E. Arbon, Y. Zhu and A. S. Mey, Journal of Chemical Theory and Computation, 2024, 20, 977–988.

58 A. V. Sinitskiy and V. S. Pande, The Journal of Chemical Physics, 2018, 148, year.

59 S.-H. Ahn, B. R. Jagger and R. E. Amaro, Journal of chemical information and modeling, 2020, 60, 5340–5352.

60 D. Aristoff, J. Copperman, G. Simpson, R. J. Webber and D. M. Zuckerman, The Journal of chemical physics, 2023, 158, year.

61 S. A. Frank, Biology direct, 2013, 8, 1–25.

62 A. Dickson and S. D. Lotz, Biophysical journal, 2017, 112, 620–629.

63 A. Bogetti, D. Yang, H. Piston, D. LeBard and L. Chong, bioRxiv, 2025, 2025–04.

64 N. M. Roussey and A. Dickson, Journal of computational chemistry, 2023, 44, 935–947.

65 W.J. Peña Ccoa and G. M. Hocky, The Journal of Chemical Physics, 2022, 156, year.

66 O. Blumer, S. Reuveni and B. Hirshberg, Journal of Chemical Theory and Computation, 2024, 20, 3484–3491.

67 M. Invernizzi and M. Parrinello, The journal of physical chemistry letters, 2020, 11, 2731– 2736.

68 D. Hamelberg, J. Mongan and J. A. McCammon, The Journal of chemical physics, 2004, 120, 11919–11929.

69 J. Wang, P. R. Arantes, A. Bhattarai, R. V. Hsu, S. Pawnikar, Y.-m. M. Huang, G. Palermo and Y. Miao, Wiley Interdisciplinary Reviews: Computational Molecular Science, 2021, 11, e1521.

70 Y. Miao, V. A. Feher and J. A. McCammon, Journal of chemical theory and computation, 2015, 11, 3584–3595.

71 A. Bhattarai and Y. Miao, Expert opinion on drug discovery, 2018, 13, 1055–1065.

72 M. M. Copeland, H. N. Do, L. Votapka, K. Joshi, J. Wang, R. E. Amaro and Y. Miao, The Journal of Physical Chemistry B, 2022, 126, 5810–5820.

73 Y. Miao, A. Bhattarai and J. Wang, Journal of chemical theory and computation, 2020, 16, 5526–5547.

74 Y. Miao, A. Bhattarai and J. Wang, Biophysical Journal, 2021, 120, 97a.

75 Y. Wang, J. Fass, B. Kaminow, J. E. Herr, D. Rufa, I. Zhang, I. Pulido, M. Henry, H. E. B. Macdonald, K. Takaba et al., Chemical Science, 2022, 13, 12016–12033.

76 K. Takaba, A. J. Friedman, C. E. Cavender, P. K. Behara, I. Pulido, M. M. Henry, H. MacDermott-Opeskin, C. R. Iacovella, A. M. Nagle, A. M. Payne et al., Chemical Science, 2024, 15, 12861–12878.

77 A. A. Ojha, L. W. Votapka, G. A. Huber, S. Gao and R. E. Amaro, Living Journal of Computational Molecular Science, 2023, 5, 2359–2359.

78 L. W. Votapka, A. M. Stokely, A. A. Ojha and R. E. Amaro, Journal of chemical information and modeling, 2022, 62, 3253–3262.

79 L. W. Votapka, B. R. Jagger, A. L. Heyneman and R. E. Amaro, The Journal of Physical Chemistry B, 2017, 121, 3597–3606.

80 B. R. Jagger, A. A. Ojha and R. E. Amaro, Journal of Chemical Theory and Computation, 2020, 16, 5348–5357.

81 L. W. Votapka and R. E. Amaro, PLoS computational biology, 2015, 11, e1004381.

82 E. Vanden-Eijnden and M. Venturoli, The Journal of chemical physics, 2009, 130, year.

83 D. Huang and A. Caflisch, PLoS computational biology, 2011, 7, e1002002.

84 S. Wolf, B. Lickert, S. Bray and G. Stock, Nature communications, 2020, 11, 2918.

85 F. Pietrucci, F. Marinelli, P. Carloni and A. Laio, Journal of the American Chemical Society, 2009, 131, 11811–11818.

86 S. Rathnayake, B. Narayan, R. Elber and C. F. Wong, Proteins: Structure, Function, and Bioinformatics, 2023, 91, 209–217.

87 G. A. Huber and J. A. McCammon, Computer Physics Communications, 2010, 181, 1896– 1905.

88 G. A. Huber and J. A. McCammon, Trends in chemistry, 2019, 1, 727–738.

89 S. Zhou, R. G. Weiß, L.-T. Cheng, J. Dzubiella, J. A. McCammon and B. Li, Proceedings of the National Academy of Sciences, 2019, 116, 14989–14994.

90 M. E. Davis, J. D. Madura, B. A. Luty and J. A. McCammon, Computer Physics Communications, 1991, 62, 187–197.

91 A. Rojnuckarin, D. R. Livesay and S. Subramaniam, Biophysical journal, 2000, 79, 686– 693.

92 J. C. Phillips, R. Braun, W. Wang, J. Gumbart, E. Tajkhorshid, E. Villa, C. Chipot, R. D. Skeel, L. Kale and K. Schulten, Journal of computational chemistry, 2005, 26, 1781–1802.

93 J. C. Phillips, D. J. Hardy, J. D. Maia, J. E. Stone, J. V. Ribeiro, R. C. Bernardi, R. Buch, G. Fiorin, J. Hénin, W. Jiang et al., The Journal of chemical physics, 2020, 153, year.

94 C. W. Hopkins, S. Le Grand, R. C. Walker and A. E. Roitberg, Journal of chemical theory and computation, 2015, 11, 1864–1874.

95 F. Sohraby and A. Nunes-Alves, Trends in Biochemical Sciences, 2023, 48, 437–449.

96 V. Limongelli, Wiley Interdisciplinary Reviews: Computational Molecular Science, 2020, 10, e1455.

97 A. A. Ojha, A. Srivastava, L. W. Votapka and R. E. Amaro, Journal of Chemical Information and Modeling, 2023, 63, 2469–2482.

98 F. Perner, C. Perner, T. Ernst and F. H. Heidel, Cells, 2019, 8, 854.

99 X. Hu, J. Li, M. Fu, X. Zhao and W. Wang, Signal transduction and targeted therapy, 2021, 6, 402.

100 T.-H. Wei, M.-Y. Lu, S.-H. Yao, Y.-Q. Hong, J. Yang, M.-Y. Zhang, Y.-Q. Yin, Y.-J. Han, Q.-Q. Li, Z.-X. Wang et al., MedComm–Oncology, 2024, 3, e69.

101 P. C. Taylor, E. Choy, X. Baraliakos, Z. Szekanecz, R. M. Xavier, J. D. Isaacs, S. Strengholt, J. M. Parmentier, R. Lippe and Y. Tanaka, Rheumatology, 2024, 63, 298–308.

102 H. Guan, M. L. Lamb, B. Peng, S. Huang, N. DeGrace, J. Read, S. Hussain, J. Wu, C. Rivard, M. Alimzhanov et al., Bioorganic & medicinal chemistry letters, 2013, 23, 3105–3110.

103 R. A. Friesner, Advances in protein chemistry, 2005, 72, 79–104.

104 J. W. Ponder and D. A. Case, Advances in protein chemistry, 2003, 66, 27–85.

105 A. A. Ojha, L. W. Votapka and R. E. Amaro, Chemical Science, 2023, 14, 13159–13175.

106 S. Izrailev, S. Stepaniants, B. Isralewitz, D. Kosztin, H. Lu, F. Molnar, W. Wriggers and K. Schulten, Computational Molecular Dynamics: Challenges, Methods, Ideas: Proceedings of the 2nd International Symposium on Algorithms for Macromolecular Modelling, Berlin, May 21–24, 1997, 1999, pp. 39–65.

107 L. W. Votapka, A. A. Ojha, N. Asada and R. E. Amaro, The Journal of Physical Chemistry Letters, 2024, 15, 10473–10478.

108 G. Bussi and A. Laio, Nature Reviews Physics, 2020, 2, 200–212.

109 B. Ensing, M. De Vivo, Z. Liu, P. Moore and M. L. Klein, Accounts of chemical research, 2006, 39, 73–81.

110 D. J. Price and C.L. Brooks III, The Journal of chemical physics, 2004, 121, 10096–10103.

111 P. Eastman, R. Galvelis, R. P. Peláez, C. R. Abreu, S. E. Farr, E. Gallicchio, A. Gorenko, M. M. Henry, F. Hu, J. Huang et al., The Journal of Physical Chemistry B, 2023, 128, 109–116.

112 Q. Du, V. Faber and M. Gunzburger, SIAM review, 1999, 41, 637–676.

113 D. B. Kokh, B. Doser, S. Richter, F. Ormersbach, X. Cheng and R. C. Wade, The Journal of chemical physics, 2020, 153, year.

114 T. J. Dolinsky, P. Czodrowski, H. Li, J. E. Nielsen, J. H. Jensen, G. Klebe and N. A. Baker, Nucleic acids research, 2007, 35, W522–W525.

115 R. J. Simpson, Cold Spring Harbor Protocols, 2006, 2006, pdb–prot4550.

116 E. Di Cera, IUBMB life, 2009, 61, 510–515.

117 M. M. Lerch and F. S. Gorelick, Medical Clinics of North America, 2000, 84, 549–563.

118 M. Hirota, M. Ohmuraya and H. Baba, Journal of gastroenterology, 2006, 41, 832–836.

119 X. Zhan, J. Wan, G. Zhang, L. Song, F. Gui, Y. Zhang, Y. Li, J. Guo, R. K. Dawra, A. K. Saluja et al., American Journal of Physiology-Gastrointestinal and Liver Physiology, 2019, 316, G816–G825.

120 D. Rauh, G. Klebe and M. T. Stubbs, Journal of molecular biology, 2004, 335, 1325–1341.

121 M. Renatus, W. Bode, R. Huber, J. Stuerzebecher and M. T. Stubbs, Journal of medicinal chemistry, 1998, 41, 5445–5456.

122 D. Jiao, P. A. Golubkov, T. A. Darden and P. Ren, Proceedings of the National Academy of Sciences, 2008, 105, 6290–6295.

123 I. Buch, T. Giorgino and G. De Fabritiis, Proceedings of the National Academy of Sciences, 2011, 108, 10184–10189.

124 A. de Ruiter and C. Oostenbrink, Journal of Chemical Theory and Computation, 2012, 8, 3686–3695.

125 D. Rauh, S. Reyda, G. Klebe and M. T. Stubbs, Trypsin mutants for structure-based drug design: expression, refolding and crystallisation, 2002.

126 F. Guillain and D. Thusius, Journal of the American Chemical Society, 1970, 92, 5534– 5536.

127 D. Mahalingam, R. Swords, J. S. Carew, S. Nawrocki, K. Bhalla and F. Giles, British journal of cancer, 2009, 100, 1523–1529.

128 M. M. Biebl and J. Buchner, Cold Spring Harbor perspectives in biology, 2019, 11, a034017.

129 A. Khandelwal, C. N. Kent, M. Balch, S. Peng, S. J. Mishra, J. Deng, V. W. Day, W. Liu, C. Subramanian, M. Cohen et al., Nature communications, 2018, 9, 425.

130 K. H. Yim, T. L. Prince, S. Qu, F. Bai, P. A. Jennings, J. N. Onuchic, E. A. Theodorakis and L. Neckers, Proceedings of the National Academy of Sciences, 2016, 113, E4801–E4809.

131 F. Chen, C. Tang, F. Yang, A. Ekpenyong, R. Qin, J. Xie, F. Momen-Heravi, N. F. Saba and Y. Teng, Science Advances, 2024, 10, eadk3663.

132 B. Ory, M. Baud’Huin, F. Verrecchia, B. B.-L. Royer, T. Quillard, J. Amiaud, S. Battaglia, D. Heymann, F. Redini and F. Lamoureux, Clinical Cancer Research, 2016, 22, 2520–2533.

133 Y. Miyata, H. Nakamoto and L. Neckers, Current pharmaceutical design, 2013, 19, 347– 365.

134 H.-K. Park, N. G. Yoon, J.-E. Lee, S. Hu, S. Yoon, S. Y. Kim, J.-H. Hong, D. Nam, Y. C. Chae, J. B. Park et al., Experimental & molecular medicine, 2020, 52, 79–91.

135 J. Beliakoff and L. Whitesell, Anti-cancer drugs, 2004, 15, 651–662.

136 A. Maloney and P. Workman, Expert opinion on biological therapy, 2002, 2, 3–24.

137 J. Trepel, M. Mollapour, G. Giaccone and L. Neckers, Nature reviews cancer, 2010, 10, 537–549.

138 N. Iqbal and N. Iqbal, Molecular biology international, 2014, 2014, 852748.

139 R. S. Herbst, International Journal of Radiation Oncology* Biology* Physics, 2004, 59, S21– S26.

140 R. Chiarle, C. Voena, C. Ambrogio, R. Piva and G. Inghirami, Nature Reviews Cancer, 2008, 8, 11–23.

141 H. Wei, Y. Zhang, Y. Jia, X. Chen, T. Niu, A. Chatterjee, P. He and G. Hou, MedComm, 2024, 5, e470.

142 J. Y. Kim, T.-M. Cho, J. M. Park, S. Park, M. Park, K. D. Nam, D. Ko, J. Seo, S. Kim, E. Jung et al., Oncogene, 2022, 41, 3289–3297.

143 M. Amaral, D. Kokh, J. Bomke, A. Wegener, H. Buchstaller, H. Eggenweiler, P. Matias, C. Sirrenberg, R. Wade and M. Frech, Nature communications, 2017, 8, 2276.

144 S. Wolf, B. Sohmen, B. Hellenkamp, J. Thurn, G. Stock and T. Hugel, bioRxiv, 2020, 2020– 02.

145 S. Wolf, M. Amaral, M. Lowinski, F. Vallée, D. Musil, J. Guldenhaupt, M. K. Dreyer, J. Bomke, M. Frech, J. Schlitter et al., Journal of Chemical Information and Modeling, 2019, 59, 5135–5147.

146 S. E. Jackson, Molecular chaperones, 2013, 155–240.

147 R. Shaknovich, G. Shue and D. Stave Kohtz, Molecular and cellular biology, 1992, 12, 5059–5068.

148 C. Antonsson, M. L. Whitelaw, J. McGuire, J.-Å. Gustafsson and L. Poellinger, Molecular and cellular biology, 1995, 15, 756–765.

149 A. Hoter, M. E. El-Sabban and H. Y. Naim, International journal of molecular sciences, 2018, 19, 2560.

150 L. H. Pearl and C. Prodromou, Annu. Rev. Biochem., 2006, 75, 271–294.

151 P. A. Brough, L. Baker, S. Bedford, K. Brown, S. Chavda, V. Chell, J. D’Alessandro, N. G. Davies, B. Davis, L. Le Strat et al., Journal of Medicinal Chemistry, 2017, 60, 2271–2286.

152 F. Vallée, C. Carrez, F. Pilorge, A. Dupuy, A. Parent, L. Bertin, F. Thompson, P. Ferrari, F. Fassy, A. Lamberton et al., Journal of medicinal chemistry, 2011, 54, 7206–7219.

153 D. A. Schuetz, L. Richter, M. Amaral, M. Grandits, U. Gradler, D. Musil, H.-P. Buchstaller, H.-M. Eggenweiler, M. Frech and G. F. Ecker, Journal of Medicinal Chemistry, 2018, 61, 4397–4411.

154 S. T. Pachis and G. J. Kops, Open biology, 2018, 8, 180109.

155 F. Althoff, R. E. Karess and C. F. Lehner, Molecular biology of the cell, 2012, 23, 2275–2291.

156 M. Schmidt, Y. Budirahardja, R. Klompmaker and R. H. Medema, EMBO reports, 2005, 6, 866–872.

157 Z. Zhou, M. He, A. A. Shah and Y. Wan, Cell division, 2016, 11, 1–18.

158 Z. Ji, H. Gao, L. Jia, B. Li and H. Yu, Elife, 2017, 6, e22513.

159 L. Sun, X. Chen, C. Song, W. Shi, L. Liu, S. Bai, X. Wang, J. Chen, C. Jiang, S.-m. Wang et al., Elife, 2024, 13, RP97896.

160 P. Lara-Gonzalez, J. Pines and A. Desai, Seminars in cell & developmental biology, 2021, pp. 86–98.

161 N. Lu and L. Ren, Bioengineered, 2021, 12, 5759–5768.

162 R. Liu, Y. Lu, J. Li, W. Yao, J. Wu, X. Chen, L. Huang, D. Nan, Y. Zhang, W. Chen et al., Cell Death & Disease, 2024, 15, 291.

163 W. Yao, M. Jiang, M. Zhang, H. Zhang and X. Liang, Journal of Cellular Signaling, 2021, 2, 212–220.

164 H. Zhang, W. Yao, M. Zhang, Y. Lu, J. Tang, M. Jiang, X. Mou, G. You and X. Liang, Biochemical and Biophysical Research Communications, 2021, 550, 84–91.

165 T. Mühlenberg, J. Falkenhorst, T. Schulz, B. S. Fletcher, A. Teuber, D. Krzeciesa, I. Klooster, M. Lundberg, L. Wilson, J. Lategahn et al., Journal of Clinical Oncology, 2024, 42, 1439– 1449.

166 A. Koch, A. Maia, A. Janssen and R. Medema, Oncogene, 2016, 35, 2518–2528.

167 J. M. Mason, X. Wei, G. C. Fletcher, R. Kiarash, R. Brokx, R. Hodgson, I. Beletskaya, M. R. Bray and T. W. Mak, Proceedings of the National Academy of Sciences, 2017, 114, 3127– 3132.

168 B. Wang, H. Wu, C. Hu, H. Wang, J. Liu, W. Wang and Q. Liu, Signal Transduction and Targeted Therapy, 2021, 6, 423.

169 S. Re, H. Oshima, K. Kasahara, M. Kamiya and Y. Sugita, Proceedings of the National Academy of Sciences, 2019, 116, 18404–18409.

170 R. Colombo, M. Caldarelli, M. Mennecozzi, M. L. Giorgini, F. Sola, P. Cappella, C. Perrera, S. R. Depaolini, L. Rusconi, U. Cucchi et al., Cancer research, 2010, 70, 10255–10264.

171 Y. Hiruma, A. Koch, S. Dharadhar, R. P. Joosten and A. Perrakis, Proteins: Structure, Function, and Bioinformatics, 2016, 84, 1761–1766.

172 J. C. Uitdehaag, J. de Man, N. Willemsen-Seegers, M. B. Prinsen, M. A. Libouban, J. G. Sterrenburg, J. J. de Wit, J. R. de Vetter, J. A. de Roos, R. C. Buijsman et al., Journal of molecular biology, 2017, 429, 2211–2230.

173 N. Kwiatkowski, N. Jelluma, P. Filippakopoulos, M. Soundararajan, M. S. Manak, M. Kwon, H. G. Choi, T. Sim, Q. L. Deveraux, S. Rottmann et al., Nature chemical biology, 2010, 6, 359–368.

174 R. Capelli, W. Lyu, V. Bolnykh, S. Meloni, J. M. H. Olsen, U. Rothlisberger, M. Parrinello and P. Carloni, The journal of physical chemistry letters, 2020, 11, 6373–6381.

175 L. A. Earl, V. Falconieri, J. L. Milne and S. Subramaniam, Current opinion in structural biology, 2017, 46, 71–78.

176 A. A. Ojha, R. Blackwell, E. R. Cruz-Chú, R. Dsouza, M. A. Astore, P. Schwander and S. M. Hanson, Biological Crystallography, 2025, 81, year.

177 H. Zheng, K. B. Handing, M. D. Zimmerman, I. G. Shabalin, S. C. Almo and W. Minor, Expert opinion on drug discovery, 2015, 10, 975–989.

178 M. Pellecchia, D. S. Sem and K. Wüthrich, Nature Reviews Drug Discovery, 2002, 1, 211– 219.

179 M. Post, S. Wolf and G. Stock, Journal of Chemical Theory and Computation, 2023, 19, 8978–8986.

180 V. Tanzel, M. Jager and S. Wolf, Journal of Chemical Theory and Computation, 2024, 20, 5058–5067.

181 S. Wolf, Journal of Chemical Information and Modeling, 2023, 63, 2902–2910.

182 B. J. Grant, A. A. Gorfe and J. A. McCammon, Current opinion in structural biology, 2010, 20, 142–147.

183 D. E. Koshland, Nature medicine, 1998, 4, 1112–1114.

184 X. Nuqui, L. Casalino, L. Zhou, M. Shehata, A. Wang, A. L. Tse, A. A. Ojha, F. L. Kearns, M. A. Rosenfeld, E. H. Miller et al., Nature Communications, 2024, 15, 7370.

