## Supplemental Information File for "seekrflow: Towards end-to-end automated simulation pipeline with machine-learned force fields for accelerated drug-target kinetic and thermodynamic predictions"

**Table S1.** Enhanced sampling approaches for estimating $k_{\text{off}}$ for receptor–ligand complexes.

| Method[a] | Receptor-ligand complex[b] | Sampling time (s) | $\tau^{\text{simulation}}$ (s) | $\tau^{\text{experiment}}$ (s) | $k_{\text{off}}^{\text{sim}}$ ($s^{-1}$) | $k_{\text{off}}^{\text{exp}}$ ($s^{-1}$) | Reference |
|---|---|---|---|---|---|---|---|
| $\text{OPES}_f$ | Trypsin-benzamidine | – | $6.40\times10^{-4}$ | $1.70\times10^{-3}$ | $1.56\times10^{3}$ | $6.00\times10^{2}$ | 1 |
| iMetaD | Trypsin-benzamidine | – | – | – | $(4.176\pm0.324)\times10^{3}$ | $6.00\times10^{2}$ | 2 |
| iMetaD | Trypsin-benzamidine | $5\times10^{-6}$ | – | – | $(9.10\pm2.50)\times10^{0}$ | $6.00\times10^{2}$ | 3 |
| LiGaMD | Trypsin-benzamidine | $5\times10^{-6}$ | – | – | $(3.53\pm1.41)\times10^{0}$ | $6.00\times10^{2}$ | 4 |
| MSM (CGMD) | Trypsin-benzamidine | $3.98\times10^{-4}$ | – | – | $6.90\times10^{5}$ | $6.00\times10^{2}$ | 5 |
| WE (REVO) | Trypsin-benzamidine | $8.75\times10^{-6}$ | $3.76\times10^{-3}$ | $1.70\times10^{-3}$ | – | – | 6 |
| WE (WExplore) | Trypsin-benzamidine | $8.75\times10^{-6}$ | $1.19\times10^{-3}$ | $1.70\times10^{-3}$ | – | – | 6 |
| WE (WExplore) | Trypsin-benzamidine | $4.10\times10^{-6}$ | $1.80\times10^{-4}$ | $1.70\times10^{-3}$ | – | – | 7 |
| dcTMD | Trypsin-benzamidine | $1\times10^{-2}$ (1D Langevin) | – | – | $(2.70\pm0.40)\times10^{2}$ | $6.00\times10^{2}$ | 8 |
| M-WEM | Trypsin-benzamidine | $4.8\times10^{-7}$ | $1.26\times10^{-3}$ | $1.70\times10^{-3}$ | $(7.91\pm1.97)\times10^{2}$ | $6.00\times10^{2}$ | 9 |
| MSM | Trypsin-benzamidine | $4.95\times10^{-5}$ | $(1.17\pm0.43)\times10^{-5}$ | $1.70\times10^{-3}$ | $(9.5\pm3.3)\times10^{4}$ | $6.00\times10^{2}$ | 10 |
| MSM | Trypsin-benzamidine | $1.49\times10^{-4}$ | – | – | $(1.31\pm1.09)\times10^{4}$ | $6.00\times10^{2}$ | 11 |
| MSM | Trypsin-benzamidine | $4.89\times10^{-5}$ | – | – | $2.80\times10^{4}$ | $6.00\times10^{2}$ | 12 |
| MSM | Trypsin-benzamidine | $5.23\times10^{-5}$ | – | – | $1.17\times10^{3}$ | $6.00\times10^{2}$ | 13 |
| AMS | Trypsin-benzamidine | $2.30\times10^{-6}$ | – | – | $(2.6\pm2.4)\times10^{2}$ | $6.00\times10^{2}$ | 14 |
| LiGaMD2 | T4L:L99A-benzene | $3\times10^{-6}$ | – | – | $(1.44\pm0.88)\times10^{3}$ | $9.50\times10^{2}$ | 15 |
| | T4L:M102A-benzene | $3\times10^{-6}$ | – | – | $(2.01\pm1.61)\times10^{3}$ | $3.00\times10^{3}$ | 15 |
| | T4L:F104A-benzene | $3\times10^{-6}$ | – | – | $(1.38\pm0.67)\times10^{6}$ | $>1.00\times10^{4}$ | 15 |
| | T4L:L99A-indole | $3\times10^{-6}$ | – | – | $(3.49\pm0.56)\times10^{3}$ | $3.25\times10^{2}$ | 15 |
| FaMetaD | T4L:L99A-benzene | $5.5\times10^{-6}$ | $(1.76\pm0.68)\times10^{-1}$ | $1.05\times10^{-3}$ | – | $9.50\times10^{2}$ | 16 |
| | T4L:L99A-indole | $2\times10^{-6}$ | $(1.68\pm0.95)\times10^{-1}$ | $3.08\times10^{-3}$ | – | $3.25\times10^{2}$ | 16 |
| iMetaD | T4L:L99A-benzene | $6.7\times10^{-6}$ | $(1.68\pm0.59)\times10^{-1}$ | $1.05\times10^{-3}$ | – | $9.50\times10^{2}$ | 16 |
| | T4L:L99A-indole | $4.5\times10^{-6}$ | $(1.02\pm0.87)\times10^{-1}$ | $3.08\times10^{-3}$ | – | $3.25\times10^{2}$ | 16 |
| iMetaD | T4L:L99A-benzene | $1.2\times10^{-5}$ | – | – | $(7.00\pm2.00)\times10^{0}$ | $(8.00\pm2.00)\times10^{2}$ | 17 |
| PIB | T4L:L99A-benzene | – | – | – | $(3.30\pm0.80)\times10^{0}$ | $(8.00\pm2.00)\times10^{2}$ | 18 |
| MSM | T4L:L99A-benzene | $6.0\times10^{-5}$ | – | – | $(3.10\pm1.30)\times10^{2}$ | $9.50\times10^{2}$ | 19 |
| iMetaD | T4L:L99A-benzene | – | – | – | $(2.70\pm1.00)\times10^{2}$ | $9.50\times10^{2}$ | 19 |
| SPIB-iMetaD | T4L:L99A-benzene | – | $1.30\times10^{-3}$ | $(1.30\pm0.30)\times10^{-3}$ | – | $(8.00\pm2.00)\times10^{2}$ | 20 |
| TS-PPTIS | c-Src kinase–imatinib | – | – | – | $2.60\times10^{-2}$ | $(1.10\pm0.80)\times10^{-1}$ | 21 |
| iMetaD | c-Src kinase–dasatinib | – | $(2.10\pm1.00)\times10^{1}$ | $1.80\times10^{1}$ | $(4.80\pm2.40)\times10^{-2}$ | $6.00\times10^{-2}$ | 22 |
| Milestoning | Abl kinase–imatinib | $\approx1\times10^{-6}$ | $5.50\times10^{-2}$ | $(4.00\pm1.00)\times10^{-2}$ | – | $(2.50\pm0.60)\times10^{1}$ | 23 |
| iMetaD | Abl kinase–imatinib | – | $(1.60\pm0.80)\times10^{3}$ | $(1.20\pm0.12)\times10^{3}$ | $(6.00\pm3.00)\times10^{-4}$ | $(8.30\pm0.83)\times10^{-4}$ | 24 |
| | Abl:N368S-imatinib | – | $(3.00\pm2.00)\times10^{2}$ | $(3.70\pm0.37)\times10^{2}$ | $(4\pm2)\times10^{-3}$ | $(2.7\pm0.27)\times10^{-3}$ | 24 |
| SPIB-iMetaD | Abl kinase-imatinib | – | $1.15\times10^{3}$ | $(1.20\pm0.12)\times10^{3}$ | – | – | 20 |
| | Abl:N368S-imatinib | – | $3.30\times10^{2}$ | $(4.00\pm0.40)\times10^{2}$ | – | – | 20 |
| | Abl:L364I-imatinib | – | $1.00\times10^{4}$ | $(7.00\pm0.70)\times10^{2}$ | – | – | 20 |
| WE(WExplore) | Epoxide hydrolase–TPPU | $6\times10^{-6}$ | $4.20\times10^{1}$ | $6.60\times10^{2}$ | – | – | 25 |
| WE (REVO) | Translocator protein–PK-11195 | $4.0\times10^{-5}$ | $1.68\times10^{3}$ | $2.04\times10^{3}$ | – | – | 26 |
| iMetaD | p38 MAP kinase–tBu-pTol-PzU | $6.80\times10^{-6}$ | – | – | $(2.0\pm1.1)\times10^{-2}$ | $1.4\times10^{-1}$ | 27 |
| faMetaD | M2 receptor–iperoxo | $8.00\times10^{-6}$ | $(2.7\pm0.5)\times10^{3}$ | – | $(3.7\pm0.7)\times10^{-4}$ | $(1.0\pm0.2)\times10^{-2}$ | 28 |
| iMetaD | $\mu$-opioid receptor–morphine | $\approx4.00\times10^{-6}$ | – | – | $(5.32\,(4.60,6.01))\times10^{-2}$ | $(2.313\pm0.167)\times10^{-2}$ | 29 |
| | $\mu$-opioid receptor–buprenorphine | $\approx1.90\times10^{-5}$ | – | – | $(2.11\,(1.40,2.70))\times10^{-2}$ | $(1.770\pm0.330)\times10^{-3}$ | 29 |
| iMetaD | $\mu$-opioid receptor–fentanyl | $\approx6.00\times10^{-6}$ | $3.80\times10^{1}$ | $\approx2.40\times10^{2}$ | – | – | 30 |
| $\tau$RAMD+extrapolation | Catalase–isoniazid | – | $3.61\times10^{1}$ | $(5.0\pm0.8)\times10^{1}$ | – | – | 31 |
| dcTMD | HSP90–resorcinol | $5\times10^{-3}$ (1D Langevin) | – | – | $(1.6\pm0.2)\times10^{-3}$ | $(3.40\pm0.20)\times10^{-2}$ | 8 |
| LiGaMD3 | MDM2–nutlin | $6.00\times10^{-6}$ | – | – | $(1.545\pm0.469)\times10^{1}$ | $4.80\times10^{-1}$ | 32 |
| LiGaMD | MDM2–nutlin | $6.00\times10^{-6}$ | – | – | $(6.951\pm5.837)\times10^{1}$ | $4.80\times10^{-1}$ | 32 |
| iMetaD | $\alpha$7-nAChR–[18F]ASEM | – | – | – | $3.30\times10^{-3}$ | $3.00\times10^{-4}$ | 33 |
| Milestoning | PYK2–ligand 1 | $5.3\times10^{-7}$ | – | – | $4.28\times10^{0}$ | $(7.40\pm2.50)\times10^{-2}$ | 34 |
| | PYK2–ligand 2 (AMBER FF) | $5.3\times10^{-7}$ | – | – | $2.06\times10^{4}$ | $(7.50\pm3.20)\times10^{-1}$ | 34 |
| | PYK2–ligand 2 (CHARMM FF) | $1.31\times10^{-6}$ | – | – | $3.03\times10^{3}$ | $(7.50\pm3.20)\times10^{-1}$ | 34 |
| | PYK2–ligand 8 | $8.6\times10^{-7}$ | – | – | $9.59\times10^{-2}$ | $(2.50\pm1.10)\times10^{-1}$ | 34 |
| Milestoning | GSK-3$\beta$ | $8.8\times10^{-7}$ | $(1.54\pm0.03)\times10^{1}$ | $1.84\times10^{1}$ | – | – | 35 |
| MSM | PYK2–ligand 1 | $6.51\times10^{-6}$ | – | – | 0.17–33.44[c] | $(7.40\pm2.50)\times10^{-2}$ | 36 |
| SMD+Bell-Evans model | $A_{2A}$ receptor–ZMA241385 | – | $2.50\times10^{1}$ | $(2.3808\pm0.4536)\times10^{3}$ | – | – | 37 |
| | $A_{2A}$ receptor–NECA | – | $8.6\times10^{-1}$ | $(1.1766\pm0.2214)\times10^{3}$ | – | – | 37 |
| SMD+Bell-Evans model | p38 MAP kinase–ligand B96 | $4.5\times10^{-6}$ | $4.32\times10^{3}$ | $1.91\times10^{4}$ | – | – | 38 |
| | p38 MAP kinase–ligand BMU | $4.5\times10^{-6}$ | $1.68\times10^{2}$ | $3.57\times10^{1}$ | – | – | 38 |
| | p38 MAP kinase–ligand SB5 | $4.5\times10^{-6}$ | $1.10\times10^{1}$ | $3.70\times10^{1}$ | – | – | 38 |
| | p38 MAP kinase–ligand SB6 | $4.5\times10^{-6}$ | $4.00\times10^{0}$ | $7.70\times10^{0}$ | – | – | 38 |
| | p38 MAP kinase–ligand SB7 | $4.5\times10^{-6}$ | $2.50\times10^{1}$ | $1.49\times10^{1}$ | – | – | 38 |
| | p38 MAP kinase–ligand BR5 | $4.5\times10^{-6}$ | $2.05\times10^{4}$ | $6.66\times10^{4}$ | – | – | 38 |
| | p38 MAP kinase–ligand BR8 | $4.5\times10^{-6}$ | $2.16\times10^{3}$ | $3.00\times10^{2}$ | – | – | 38 |
| | p38 MAP kinase–ligand B12 | $4.5\times10^{-6}$ | $5.00\times10^{4}$ | $3.82\times10^{4}$ | – | – | 38 |
| SMD+Bell model | Streptavidin–biotin | $1.3\times10^{-5}$ | $8.73\times10^{-5}$ | $9.52\times10^{2}$ | $1.14\times10^{4}$ | $1.05\times10^{-3}$ | 39 |
| SMD+DHS model | Streptavidin–biotin | $1.3\times10^{-5}$ | $2.41\times10^{7}$ | $9.52\times10^{2}$ | $4.16\times10^{-8}$ | $1.05\times10^{-3}$ | 39 |
| iMetaD+KTR | YCD–5-fluorocytosine | – | – | – | $4.0\times10^{1}$ | $3.1\times10^{1}$ | 40 |
| iMetaD+KTR | CDK2–ligand CS3 | $1.5\times10^{-5}$ | – | – | $\log(k_{\text{off}})\approx\log(8.4)\pm\log(5.1)$ | $2.59\times10^{-1}$ | 41 |

[a] **Method abbreviations:** $\text{OPES}_f$ = On-the-fly Probability Enhanced Sampling flooding; iMetaD = Infrequent Metadynamics; LiGaMD = Ligand Gaussian-accelerated Molecular Dynamics; MSM = Markov State Model; CGMD = Coarse-grained Molecular Dynamics; WE = Weighted Ensemble; REVO = Resampling of Ensembles by Variation Optimization; dcTMD = Dissipation-corrected Targeted Molecular Dynamics; M-WEM = Markovian Weighted Ensemble Milestoning; AMS = Adaptive Multilevel Splitting; FaMetaD = frequency-adaptive Metadynamics; PIB = Predictive Information Bottleneck; SPIB-iMetaD = State Predictive Information Bottleneck Infrequent Metadynamics; TS-PPTIS = Transition State Partial Path Transition Interface Sampling; $\tau$RAMD = $\tau$-Random Acceleration Molecular Dynamics; SMD = Steered Molecular Dynamics; DHS model = Dudko-Hummer-Szabo model; KTR = Kramers' Time-dependent Rate theory

[b] **Receptor-ligand complex abbreviations:** T4L = T4 Lysozyme; c-Src = Cellular Src; TPPU = N-(1-(1-oxopropyl)-2-pyrrolidinyl)-N′-(4-trifluoromethylphenyl)urea; MAP = Mitogen-Activated Protein; tBu-pTol-PzU = 1-(3-(tert-butyl)-1-(p-tolyl)-1H-pyrazol-5-yl)urea; M2 = Muscarinic M2; HSP90 = Heat Shock Protein 90; MDM2 = Murine Double Minute 2; $\alpha$7-nAChR = $\alpha$7-nicotinic acetylcholine receptor; [18F]ASEM = 3-(1,4-diazabicyclo[3.2.2]nonan-4-yl)-6-[18F]fluorodibenzo[b,d]thiophene 5,5-dioxide; PYK2 = Proline-Rich Tyrosine Kinase 2; GSK-3$\beta$ = Glycogen Synthase Kinase-3$\beta$; $A_{2A}$ = Adenosine type 2 A; YCD = Yeast Cytosine Deaminase; CDK2 = Cyclin-Dependent Kinase 2; CS3 = *S*-[3-oxo-3-(2-thienyl)propyl]-L-cysteine

[c] The manuscript presents a range of values, depending on the MSM parameters.

**Table S2.** Milestoning theory for receptor-ligand kinetics and thermodynamics[42,43,44].

| Equation | Description |
|---|---|
| $\mathbf{Q} = \begin{pmatrix} q_{1,1} & q_{1,2} & \cdots & q_{1,N} \\ q_{2,1} & q_{2,2} & \cdots & q_{2,N} \\ \vdots & \vdots & \ddots & \vdots \\ q_{N,1} & q_{N,2} & \cdots & q_{N,N} \end{pmatrix}$ | $\mathbf{Q}$ - $N \times N$ transition rate matrix<br>$q_{ij}$ - Transition rate from milestone $i$ to $j$ |
| $q_{ii} = -\sum_{j \neq i} q_{ij}$ | Diagonal elements - Self-transition rate |
| $q_{ij} = \begin{cases} \frac{N_{ij}}{R_i} & \text{if } R_i \neq 0 \\ 0 & \text{if } R_i = 0 \end{cases}$ | $N_{ij}$ - Number of transitions between $i$ to $j$<br>$R_i$ - Residence time after last contact with milestone $i$ |
| $x_\alpha(t+\Delta t) = \begin{cases} x_\alpha{}^* & \text{if } x_\alpha{}^* \in V_\alpha \\ x_\alpha(t) & \text{otherwise} \end{cases}$ | $x_\alpha^*$ - Predicted position at time, $t+\Delta t$<br>$V_\alpha$ - Voronoi cell, $\alpha$ |
| $v_\alpha(t+\Delta t) = \begin{cases} v_\alpha{}^* & \text{if } x_\alpha{}^* \in V_\alpha \\ -v_\alpha(t) & \text{otherwise} \end{cases}$ | $v_\alpha^*$ - Predicted velocity at time, $t+\Delta t$<br>-$v_\alpha^*$ - Velocity reversal if $x_\alpha^*$ is outside $V_\alpha$ |
| $T = \left( \sum_{\alpha=1}^{n} \frac{\pi_\alpha}{T_\alpha} \right)^{-1}$ | $T$ - Normalizing factor<br>$\pi_\alpha$ - Equilibrium probability of $V_\alpha$<br>$T_\alpha$ - Simulation time in $V_\alpha$ |
| $N_{ij} = T \sum_{\alpha=1}^{n} \pi_\alpha \frac{N_{ij}^\alpha}{T_\alpha}$ | $N_{ij}^\alpha$ - Collisions from milestone $j$ to $i$ in $V_\alpha$<br>$\sum_{\alpha=1}^{n}$ - Aggregated over cells weighted by $\pi_\alpha$ |
| $R_i = T \sum_{\alpha=1}^{n} \pi_\alpha \frac{R_i^\alpha}{T_\alpha}$ | $R_i^\alpha$ - Time spent in $V_\alpha$ after last touching milestone $i$<br>Total residence time at milestone $i$ |
| $\sum_{\beta \neq \alpha} \pi_\beta \frac{N_{\beta,\alpha}}{T_\beta} = \sum_{\beta \neq \alpha} \pi_\alpha \frac{N_{\alpha,\beta}}{T_\alpha}$ | Constraint 1 - Flux equilibrium |
| $\sum_{\alpha=1}^{n} \pi_\alpha = 1$ | Constraint 2 - Ensures total probability equals 1 |
| Mean free passage time (MFPT)<br>$\hat{\mathbf{Q}}\mathbf{T}^N = -\mathbf{1}$ | $\hat{\mathbf{Q}}$ - $(N-1) \times (N-1)$ submatrix<br>$\mathbf{T}^N$ - MFPT vector<br>$\mathbf{1}$ - Vector of ones |
| $\mathbf{Qp} = \mathbf{p}$ | $\mathbf{p}$ - Steady-state probability vector |
| Free energy per milestone<br>$\Delta G_i = -RT \ln \left( \frac{p_i}{p_{\text{ref}}} \right)$ | $R$ - Gas constant<br>$T$ - Temperature<br>$p_i$ and $p_{\text{ref}}$ - Stationary probabilities |

**Table S3.** Anchor and milestone positions for the trypsin-benzamidine complex.

| Complex | Anchors (nm) | Milestones (nm) |
|---|---|---|
| Trypsin-benzamidine | [0.1, 0.2, 0.3, 0.4, 0.5, 0.6, 0.7, 0.8, 0.9, 1.0, 1.1, 1.2, 1.3, 1.4, 1.5, 1.6, 1.8] | [0.15, 0.25, 0.35, 0.45, 0.55, 0.65, 0.75, 0.85, 0.95, 1.05, 1.15, 1.25, 1.35, 1.45, 1.55, 1.7] |

**Table S4.** Theoretical and experimental residence times for trypsin-benzamidine complex.

| Complex | Theoretical residence time (s) | Experimental residence time (s) |
|---|---|---|
| Trypsin-benzamidine | $(3.1 \pm 1.8) \times 10^{-3}$ | $(1.67 \pm 0.83) \times 10^{-3}$ |

**Table S5.** Theoretical and experimental $k_{\text{off}}$ rates for trypsin-benzamidine complex.

| Complex | Theoretical $k_{\text{off}}$ ($s^{-1}$) | Experimental $k_{\text{off}}$ ($s^{-1}$) |
|---|---|---|
| Trypsin-benzamidine | $(3.2 \pm 1.1) \times 10^{2}$ | $(6.0 \pm 3.0) \times 10^{2}$ |

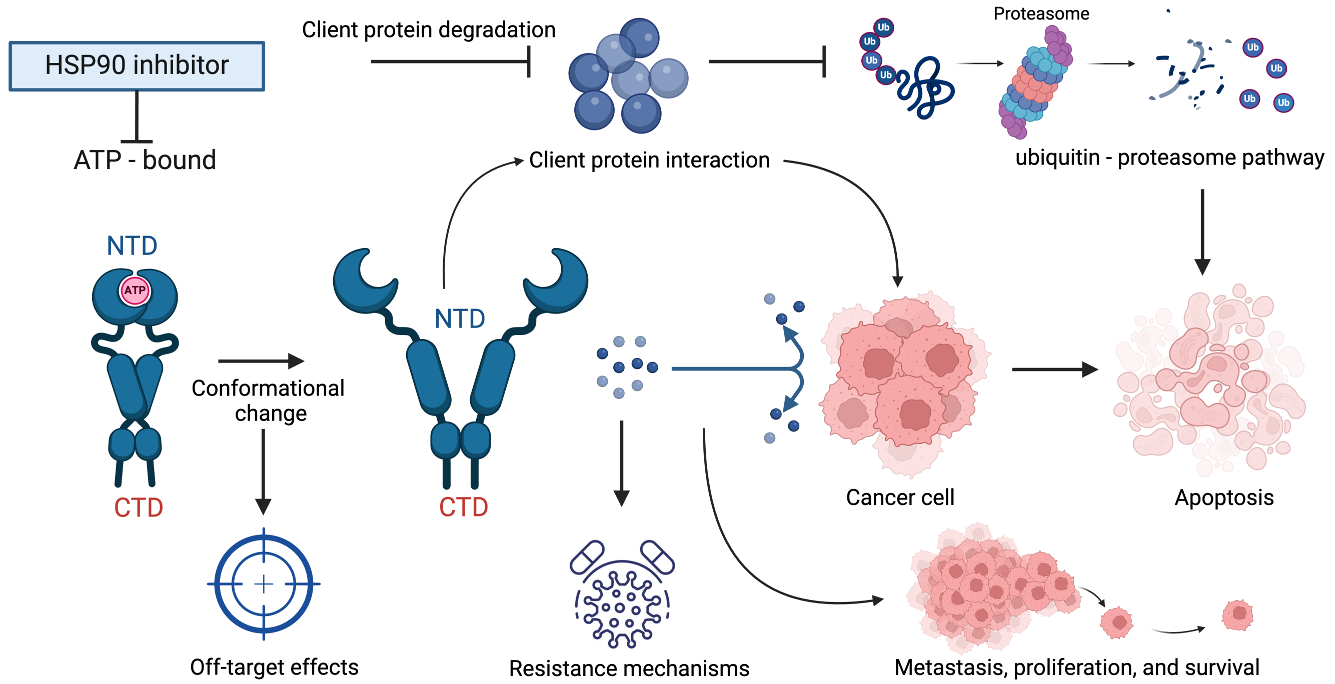


**Fig. S1** Mechanism of HSP90 inhibition and its therapeutic implications. HSP90 inhibitors target the ATP-binding pocket in the N-terminal domain (NTD) of HSP90, preventing ATP hydrolysis and disrupting the chaperone function. This leads to the degradation of client proteins via the ubiquitin-proteasome pathway, ultimately inducing apoptosis in cancer cells. By destabilizing oncogenic client proteins, HSP90 inhibitors inhibit metastasis, proliferation, and survival of tumor cells. However, resistance mechanisms and off-target effects remain challenges in developing effective HSP90-targeted therapies[45,46].

**Table S6.** HSP90-inhibitors with their respective group and binding conformation with HSP90 protein.

| Inhibitor | Group | HSP90 binding conformation |
|---|---|---|
| Inhibitor 1 | resorcinol | Helix |
| Inhibitor 2 | aminothienopyridine | Loop |
| Inhibitor 3 | aminoquinazoline | Helix |
| Inhibitor 4 | aminoquinazoline | Helix |
| Inhibitor 5 | hydroxy-indazole | Helix |
| Inhibitor 6 | 2-aminopyramidine | Helix |
| Inhibitor 7 | hydroxy-indazole | Helix |
| Inhibitor 8 | resorcinol | Loop |

**Table S7.** Theoretical and experimental $k_{off}$ rates for HSP90-inhibitor complexes.

| Complex | Theoretical $k_{off}$ ($s^{-1}$) | Experimental $k_{off}$ ($s^{-1}$) |
|---|---|---|
| HSP90-inhibitor 1 | $(2.86 \pm 0.40) \times 10^{-1}$ | $(1.10 \pm 0.43) \times 10^{-1}$ |
| HSP90-inhibitor 2 | $(1.36 \pm 0.047) \times 10^{-1}$ | $(1.09 \pm 0.19) \times 10^{-1}$ |
| HSP90-inhibitor 3 | $(5.93 \pm 0.43) \times 10^{-3}$ | $(4.54 \pm 0.15) \times 10^{-3}$ |
| HSP90-inhibitor 4 | $(4.47 \pm 0.91) \times 10^{-3}$ | $(4.53 \pm 0.41) \times 10^{-3}$ |
| HSP90-inhibitor 5 | $(2.23 \pm 0.073) \times 10^{-3}$ | $(2.01 \pm 0.20) \times 10^{-3}$ |
| HSP90-inhibitor 6 | $(2.32 \pm 0.024) \times 10^{-4}$ | $(9.89 \pm 1.30) \times 10^{-4}$ |
| HSP90-inhibitor 7 | $(1.05 \pm 0.042) \times 10^{-4}$ | $(6.79 \pm 0.053) \times 10^{-4}$ |
| HSP90-inhibitor 8 | $(1.12 \pm 0.060) \times 10^{-5}$ | $< 1.00 \times 10^{-4}$ |

**Table S8.** Theoretical and experimental residence times for HSP90-inhibitor complexes.

| Complex | Theoretical residence time (s) | Experimental residence time (s) |
|---|---|---|
| HSP90-inhibitor 1 | $(3.50 \pm 0.66) \times 10^{0}$ | $(9.09 \pm 3.55) \times 10^{0}$ |
| HSP90-inhibitor 2 | $(7.33 \pm 0.24) \times 10^{0}$ | $(9.17 \pm 1.60) \times 10^{0}$ |
| HSP90-inhibitor 3 | $(1.69 \pm 0.10) \times 10^{2}$ | $(2.20 \pm 0.07) \times 10^{2}$ |
| HSP90-inhibitor 4 | $(2.24 \pm 0.49) \times 10^{2}$ | $(2.21 \pm 0.20) \times 10^{2}$ |
| HSP90-inhibitor 5 | $(4.48 \pm 0.15) \times 10^{2}$ | $(4.98 \pm 0.50) \times 10^{2}$ |
| HSP90-inhibitor 6 | $(4.32 \pm 4.20) \times 10^{3}$ | $(1.01 \pm 0.13) \times 10^{3}$ |
| HSP90-inhibitor 7 | $(9.55 \pm 4.10) \times 10^{3}$ | $(1.47 \pm 0.01) \times 10^{3}$ |
| HSP90-inhibitor 8 | $(8.97 \pm 5.20) \times 10^{4}$ | $> 1.00 \times 10^{4}$ |

**Table S9.** Anchor and milestone positions for HSP90-inhibitor complexes.

| Complex | Anchors (nm) | Milestones (nm) |
|---|---|---|
| HSP90-inhibitor 1 | [0.25, 0.325, 0.4, 0.475, 0.55, 0.625, 0.7, 0.8, 0.9, 1.0, 1.1, 1.2, 1.3, 1.4, 1.6, 1.8, 2.0] | [0.2875, 0.3625, 0.4375, 0.5125, 0.5875, 0.6625, 0.75, 0.85, 0.95, 1.05, 1.15, 1.25, 1.35, 1.5, 1.7, 1.9] |
| HSP90-inhibitor 2 | [0.275, 0.35, 0.425, 0.5, 0.575, 0.65, 0.725, 0.8, 0.9, 1.0, 1.1, 1.2, 1.3, 1.4, 1.6, 1.8, 2.0] | [0.3125, 0.3875, 0.4625, 0.5375, 0.6125, 0.6875, 0.7625, 0.85, 0.95, 1.05, 1.15, 1.25, 1.35, 1.5, 1.7, 1.9] |
| HSP90-inhibitor 3 | [0.175, 0.25, 0.325, 0.4, 0.475, 0.55, 0.625, 0.7, 0.8, 0.9, 1.0, 1.1, 1.2, 1.4, 1.6, 1.8, 2.0] | [0.2125, 0.2875, 0.3625, 0.4375, 0.5125, 0.5875, 0.6625, 0.75, 0.85, 0.95, 1.05, 1.15, 1.3, 1.5, 1.7, 1.9] |
| HSP90-inhibitor 4 | [0.15, 0.225, 0.3, 0.375, 0.45, 0.525, 0.6, 0.7, 0.8, 0.9, 1.0, 1.1, 1.2, 1.4, 1.6, 1.8, 2.0] | [0.1875, 0.2625, 0.3375, 0.4125, 0.4875, 0.5625, 0.65, 0.75, 0.85, 0.95, 1.05, 1.15, 1.3, 1.5, 1.7, 1.9] |
| HSP90-inhibitor 5 | [0.175, 0.25, 0.325, 0.4, 0.475, 0.55, 0.625, 0.7, 0.8, 0.9, 1.0, 1.1, 1.2, 1.4, 1.6, 1.8, 2.0] | [0.2125, 0.2875, 0.3625, 0.4375, 0.5125, 0.5875, 0.6625, 0.75, 0.85, 0.95, 1.05, 1.15, 1.3, 1.5, 1.7, 1.9] |
| HSP90-inhibitor 6 | [0.1, 0.175, 0.25, 0.325, 0.4, 0.475, 0.55, 0.625, 0.7, 0.8, 0.9, 1.0, 1.2, 1.4, 1.6, 1.8, 2.0] | [0.1375, 0.2125, 0.2875, 0.3625, 0.4375, 0.5125, 0.5875, 0.6625, 0.75, 0.85, 0.95, 1.1, 1.3, 1.5, 1.7, 1.9] |
| HSP90-inhibitor 7 | [0.15, 0.225, 0.3, 0.375, 0.45, 0.525, 0.6, 0.675, 0.75, 0.825, 0.9, 1.0, 1.2, 1.4, 1.6, 1.8, 2.0] | [0.1875, 0.2625, 0.3375, 0.4125, 0.4875, 0.5625, 0.6375, 0.7125, 0.7875, 0.8625, 0.95, 1.1, 1.3, 1.5, 1.7, 1.9] |
| HSP90-inhibitor 8 | [0.3, 0.375, 0.45, 0.525, 0.6, 0.675, 0.75, 0.825, 0.9, 1.0, 1.1, 1.2, 1.3, 1.4, 1.6, 1.8, 2.0] | [0.3375, 0.4125, 0.4875, 0.5625, 0.6375, 0.7125, 0.7875, 0.8625, 0.95, 1.05, 1.15, 1.25, 1.35, 1.5, 1.7, 1.9] |

**Table S10.** Anchor and milestone positions for TTK-inhibitor complexes.

| Complex | Anchors (nm) | Milestones (nm) |
|---|---|---|
| TTK-inhibitor 1 | [0.225, 0.3, 0.375, 0.45, 0.525, 0.6, 0.7, 0.775, 0.825, 0.9, 1.0, 1.1, 1.2, 1.4, 1.6, 1.8, 2.0] | [0.2625, 0.3375, 0.4125, 0.4875, 0.5625, 0.65, 0.7375, 0.8, 0.8625, 0.95, 1.05, 1.15, 1.3, 1.5, 1.7, 1.9] |
| TTK-inhibitor 2 | [0.25, 0.325, 0.4, 0.475, 0.55, 0.625, 0.7, 0.8, 0.9, 1.0, 1.1, 1.2, 1.3, 1.4, 1.6, 1.8, 2.0] | [0.2875, 0.3625, 0.4375, 0.5125, 0.5875, 0.6625, 0.75, 0.85, 0.95, 1.05, 1.15, 1.25, 1.35, 1.5, 1.7, 1.9] |
| TTK-inhibitor 3 | [0.2, 0.275, 0.35, 0.425, 0.5, 0.575, 0.65, 0.725, 0.8, 0.9, 1.0, 1.1, 1.2, 1.4, 1.6, 1.8, 2.0] | [0.2375, 0.3125, 0.3875, 0.4625, 0.5375, 0.6125, 0.6875, 0.7625, 0.85, 0.95, 1.05, 1.15, 1.3, 1.5, 1.7, 1.9] |
| TTK-inhibitor 4 | [0.325, 0.4, 0.475, 0.55, 0.625, 0.7, 0.775, 0.85, 0.925, 1.0, 1.1, 1.2, 1.3, 1.4, 1.6, 1.8, 2.0] | [0.3625, 0.4375, 0.5125, 0.5875, 0.6625, 0.7375, 0.8125, 0.8875, 0.9625, 1.05, 1.15, 1.25, 1.35, 1.5, 1.7, 1.9] |
| TTK-inhibitor 5 | [0.25, 0.3, 0.375, 0.45, 0.525, 0.6, 0.7, 0.8, 0.9, 1.0, 1.1, 1.2, 1.3, 1.4, 1.6, 1.8, 2.0] | [0.275, 0.3375, 0.4125, 0.4875, 0.5625, 0.65, 0.75, 0.85, 0.95, 1.05, 1.15, 1.25, 1.35, 1.5, 1.7, 1.9] |
| TTK-inhibitor 6 | [0.375, 0.45, 0.525, 0.6, 0.675, 0.75, 0.825, 0.9, 1.0, 1.1, 1.2, 1.3, 1.4, 1.5, 1.6, 1.8, 2.0] | [0.4125, 0.4875, 0.5625, 0.6375, 0.7125, 0.7875, 0.8625, 0.95, 1.05, 1.15, 1.25, 1.35, 1.45, 1.55, 1.7, 1.9] |
| TTK-inhibitor 7 | [0.25, 0.3, 0.375, 0.45, 0.525, 0.6, 0.675, 0.75, 0.825, 0.9, 1.0, 1.1, 1.2, 1.4, 1.6, 1.8, 2.0] | [0.275, 0.3375, 0.4125, 0.4875, 0.5625, 0.6375, 0.7125, 0.7875, 0.8625, 0.95, 1.05, 1.15, 1.3, 1.5, 1.7, 1.9] |
| TTK-inhibitor 8 | [0.25, 0.325, 0.4, 0.475, 0.55, 0.625, 0.7, 0.8, 0.9, 1.0, 1.1, 1.2, 1.3, 1.4, 1.6, 1.8, 2.0] | [0.2875, 0.3625, 0.4375, 0.5125, 0.5875, 0.6625, 0.75, 0.85, 0.95, 1.05, 1.15, 1.25, 1.35, 1.5, 1.7, 1.9] |

**Table S11.** Theoretical and experimental $k_{off}$ rates for TTK-inhibitor complexes.

| Complex | Theoretical $k_{off}$ ($s^{-1}$) | Experimental $k_{off}$ ($s^{-1}$) |
|---|---|---|
| TTK-inhibitor 1 | $(5.72 \pm 0.12) \times 10^{0}$ | $(5.00 \pm 0.00) \times 10^{-2}$ |
| TTK-inhibitor 2 | $(9.20 \pm 1.20) \times 10^{-2}$ | $(2.80 \pm 0.00) \times 10^{-2}$ |
| TTK-inhibitor 3 | $(3.25 \pm 0.094) \times 10^{-2}$ | $(4.70 \pm 0.00) \times 10^{-2}$ |
| TTK-inhibitor 4 | $(2.08 \pm 0.28) \times 10^{-3}$ | $(8.33 \pm 0.00) \times 10^{-3}$ |
| TTK-inhibitor 5 | $(2.10 \pm 0.087) \times 10^{-3}$ | $(3.40 \pm 0.00) \times 10^{-3}$ |
| TTK-inhibitor 6 | $(3.35 \pm 0.019) \times 10^{-4}$ | $(8.33 \pm 0.00) \times 10^{-4}$ |
| TTK-inhibitor 7 | $(1.53 \pm 0.099) \times 10^{-5}$ | $(4.10 \pm 0.00) \times 10^{-4}$ |
| TTK-inhibitor 8 | $(1.42 \pm 0.099) \times 10^{-4}$ | $(1.00 \pm 0.00) \times 10^{-4}$ |

**Table S12.** Theoretical and experimental residence times for TTK-inhibitor complexes.

| Complex | Theoretical residence time (s) | Experimental residence time (s) |
|---|---|---|
| TTK-inhibitor 1 | $(1.75 \pm 0.0038) \times 10^{-1}$ | $(2.00 \pm 0.00) \times 10^{1}$ |
| TTK-inhibitor 2 | $(1.09 \pm 0.15) \times 10^{1}$ | $(3.57 \pm 0.00) \times 10^{1}$ |
| TTK-inhibitor 3 | $(3.08 \pm 0.85) \times 10^{1}$ | $(3.57 \pm 0.00) \times 10^{1}$ |
| TTK-inhibitor 4 | $(4.81 \pm 0.66) \times 10^{2}$ | $(1.20 \pm 0.00) \times 10^{2}$ |
| TTK-inhibitor 5 | $(4.77 \pm 0.20) \times 10^{2}$ | $(2.94 \pm 0.00) \times 10^{2}$ |
| TTK-inhibitor 6 | $(2.99 \pm 0.14) \times 10^{3}$ | $(1.20 \pm 0.00) \times 10^{3}$ |
| TTK-inhibitor 7 | $(6.53 \pm 0.43) \times 10^{4}$ | $(2.44 \pm 0.00) \times 10^{3}$ |
| TTK-inhibitor 8 | $(7.04 \pm 0.47) \times 10^{3}$ | $(1.00 \pm 0.00) \times 10^{4}$ |

### References

1 Narjes Ansari, Valerio Rizzi, and Michele Parrinello. Water regulates the residence time of Benzamidine in Trypsin. *Nature Communications*, 13(1):5438, September 2022. ISSN 2041-1723. doi: 10.1038/s41467-022-33104-3.

2 Z. Faidon Brotzakis, Vittorio Limongelli, and Michele Parrinello. Accelerating the Calculation of Protein–Ligand Binding Free Energy and Residence Times Using Dynamically Optimized Collective Variables. *Journal of Chemical Theory and Computation*, 15(1):743–750, January 2019. ISSN 1549-9618. doi: 10.1021/acs.jctc.8b00934.

3 Pratyush Tiwary, Vittorio Limongelli, Matteo Salvalaglio, and Michele Parrinello. Kinetics of protein–ligand unbinding: Predicting pathways, rates, and rate-limiting steps. *Proceedings of the National Academy of Sciences*, 112(5):E386–E391, February 2015. doi: 10.1073/pnas.1424461112.

4 Yinglong Miao, Apurba Bhattarai, and Jinan Wang. Ligand Gaussian Accelerated Molecular Dynamics (LiGaMD): Characterization of Ligand Binding Thermodynamics and Kinetics. *Journal of Chemical Theory and Computation*, 16(9):5526–5547, September 2020. ISSN 1549-9618. doi: 10.1021/acs.jctc.0c00395.

5 Bhupendra R. Dandekar and Jagannath Mondal. Capturing Protein–Ligand Recognition Pathways in Coarse-Grained Simulation. *The Journal of Physical Chemistry Letters*, 11(13):5302–5311, July 2020. doi: 10.1021/acs.jpclett.0c01683.

6 Nazanin Donyapour, Nicole M. Roussey, and Alex Dickson. REVO: Resampling of ensembles by variation optimization. *The Journal of Chemical Physics*, 150(24):244112, June 2019. ISSN 0021-9606. doi: 10.1063/1.5100521.

7 Alex Dickson and Samuel D. Lotz. Multiple Ligand Unbinding Pathways and Ligand-Induced Destabilization Revealed by WExplore. *Biophysical Journal*, 112:620–629, February 2017. ISSN 0006-3495. doi: 10.1016/j.bpj.2017.01.006.

8 Steffen Wolf, Benjamin Lickert, Simon Bray, and Gerhard Stock. Multisecond ligand dissociation dynamics from atomistic simulations. *Nature Communications*, 11(1):2918, June 2020. ISSN 2041-1723. doi: 10.1038/s41467-020-16655-1.

9 Dhiman Ray, Sharon Emily Stone, and Ioan Andricioaei. Markovian Weighted Ensemble Milestoning (M-WEM): Long-Time Kinetics from Short Trajectories. *Journal of Chemical Theory and Computation*, 18(1):79–95, January 2022. ISSN 1549-9618. doi: 10.1021/acs.jctc.1c00803.

10 Ignasi Buch, Toni Giorgino, and Gianni De Fabritiis. Complete reconstruction of an enzyme-inhibitor binding process by molecular dynamics simulations. *Proceedings of the National Academy of Sciences*, 108(25):10184–10189, June 2011. doi: 10.1073/pnas.1103547108.

11 Nuria Plattner and Frank Noé. Protein conformational plasticity and complex ligand-binding kinetics explored by atomistic simulations and Markov models. *Nature Communications*, 6(1):7653, July 2015. ISSN 2041-1723. doi: 10.1038/ncomms8653.

12 S. Doerr and G. De Fabritiis. On-the-Fly Learning and Sampling of Ligand Binding by High-Throughput Molecular Simulations. *Journal of Chemical Theory and Computation*, 10(5):2064–2069, May 2014. ISSN 1549-9618. doi: 10.1021/ct400919u.

13 Hao Wu, Fabian Paul, Christoph Wehmeyer, and Frank Noé. Multiensemble Markov models of molecular thermodynamics and kinetics. *Proceedings of the National Academy of Sciences*, 113(23):E3221–E3230, June 2016. doi: 10.1073/pnas.1525092113.

14 Ivan Teo, Christopher G. Mayne, Klaus Schulten, and Tony Lelièvre. Adaptive Multilevel Splitting Method for Molecular Dynamics Calculation of Benzamidine-Trypsin Dissociation Time. *Journal of Chemical Theory and Computation*, 12(6):2983–2989, June 2016. ISSN 1549-9618. doi: 10.1021/acs.jctc.6b00277.

15 Jinan Wang and Yinglong Miao. Ligand Gaussian Accelerated Molecular Dynamics 2 (LiGaMD2): Improved Calculations of Ligand Binding Thermodynamics and Kinetics with Closed Protein Pocket. *Journal of Chemical Theory and Computation*, 19(3):733–745, February 2023. ISSN 1549-9618. doi: 10.1021/acs.jctc.2c01194.

16 Yong Wang, Omar Valsson, Pratyush Tiwary, Michele Parrinello, and Kresten Lindorff-Larsen. Frequency adaptive metadynamics for the calculation of rare-event kinetics. *The Journal of Chemical Physics*, 149(7):072309, May 2018. ISSN 0021-9606. doi: 10.1063/1.5024679.

17 Yong Wang, João Miguel Martins, and Kresten Lindorff-Larsen. Biomolecular conformational changes and ligand binding: from kinetics to thermodynamics. *Chemical Science*, 8(9):6466–6473, August 2017. ISSN 2041-6539. doi: 10.1039/C7SC01627A.

18 Yihang Wang, João Marcelo Lamim Ribeiro, and Pratyush Tiwary. Past–future information bottleneck for sampling molecular reaction coordinate simultaneously with thermodynamics and kinetics. *Nature Communications*, 10(1):3573, August 2019. ISSN 2041-1723. doi: 10.1038/s41467-019-11405-4.

19 Jagannath Mondal, Navjeet Ahalawat, Subhendu Pandit, Lewis E. Kay, and Pramodh Vallurupalli. Atomic resolution mechanism of ligand binding to a solvent inaccessible cavity in T4 lysozyme. *PLOS Computational Biology*, 14(5):e1006180, May 2018. ISSN 1553-7358. doi: 10.1371/journal.pcbi.1006180.

20 Suemin Lee, Dedi Wang, Markus A. Seeliger, and Pratyush Tiwary. Calculating Protein–Ligand Residence Times through State Predictive Information Bottleneck Based Enhanced Sampling. *Journal of Chemical Theory and Computation*, 20(14):6341–6349, July 2024. ISSN 1549-9618. doi: 10.1021/acs.jctc.4c00503.

21 Susanta Haldar, Federico Comitani, Giorgio Saladino, Christopher Woods, Marc W. van der Kamp, Adrian J. Mulholland, and Francesco Luigi Gervasio. A Multiscale Simulation Approach to Modeling Drug–Protein Binding Kinetics. *Journal of Chemical Theory and Computation*, 14(11):6093–6101, November 2018. ISSN 1549-9618. doi: 10.1021/acs.jctc.8b00687.

22 Pratyush Tiwary, Jagannath Mondal, and B. J. Berne. How and when does an anticancer drug leave its binding site? *Science Advances*, 3(5):e1700014, May 2017. doi: 10.1126/sciadv.1700014.

23 Brajesh Narayan, Nicolae-Viorel Buchete, and Ron Elber. Computer Simulations of the Dissociation Mechanism of Gleevec from Abl Kinase with Milestoning. *The Journal of Physical Chemistry B*, 125(22):5706–5715, June 2021. ISSN 1520-6106. doi: 10.1021/acs.jpcb.1c00264.

24 Mrinal Shekhar, Zachary Smith, Markus A. Seeliger, and Pratyush Tiwary. Protein Flexibility and Dissociation Pathway Differentiation Can Explain Onset of Resistance Mutations in Kinases. *Angewandte Chemie International Edition*, 61(28):e202200983, 2022. ISSN 1521-3773. doi: 10.1002/anie.202200983.

25 Samuel D Lotz and Alex Dickson. Unbiased Molecular Dynamics of 11 min Timescale Drug Unbinding Reveals Transition State Stabilizing Interactions. *Journal of the American Chemical Society*, 140(2):618–628, January 2018. ISSN 0002-7863. doi: 10.1021/jacs.7b08572.

26 Tom Dixon, Arzu Uyar, Shelagh Ferguson-Miller, and Alex Dickson. Membrane-Mediated Ligand Unbinding of the PK-11195 Ligand from TSPO. *Biophysical Journal*, 120(1):158–167, January 2021. ISSN 0006-3495. doi: 10.1016/j.bpj.2020.11.015.

27 Rodrigo Casasnovas, Vittorio Limongelli, Pratyush Tiwary, Paolo Carloni, and Michele Parrinello. Unbinding Kinetics of a p38 MAP Kinase Type II Inhibitor from Metadynamics Simulations. *Journal of the American Chemical Society*, 139(13):4780–4788, April 2017. ISSN 0002-7863. doi: 10.1021/jacs.6b12950.

28 Riccardo Capelli, Wenping Lyu, Viacheslav Bolnykh, Simone Meloni, Jógvan Magnus Haugaard Olsen, Ursula Rothlisberger, Michele Parrinello, and Paolo Carloni. Accuracy of Molecular Simulation-Based Predictions of koff Values: A Metadynamics Study. *The Journal of Physical Chemistry Letters*, 11(15):6373–6381, August 2020. doi: 10.1021/acs.jpclett.0c00999.

29 João Marcelo Lamim Ribeiro, Davide Provasi, and Marta Filizola. A combination of machine learning and infrequent metadynamics to efficiently predict kinetic rates, transition states, and molecular determinants of drug dissociation from G protein-coupled receptors. *The Journal of Chemical Physics*, 153(12):124105, September 2020. ISSN 0021-9606. doi: 10.1063/5.0019100.

30 Paween Mahinthichaichan, Quynh N. Vo, Christopher R. Ellis, and Jana Shen. Kinetics and Mechanism of Fentanyl Dissociation from the -Opioid Receptor. *JACS Au*, 1(12):2208–2215, December 2021. doi: 10.1021/jacsau.1c00341.

31 Ekaterina Maximova, Eugene B. Postnikov, Anastasia I. Lavrova, Vladimir Farafonov, and Dmitry Nerukh. Protein–Ligand Dissociation Rate Constant from All-Atom Simulation. *The Journal of Physical Chemistry Letters*, 12(43):10631–10636, November 2021. doi: 10.1021/acs.jpclett.1c02952.

32 Jinan Wang and Yinglong Miao. Ligand Gaussian Accelerated Molecular Dynamics 3 (LiGaMD3): Improved Calculations of Binding Thermodynamics and Kinetics of Both Small Molecules and Flexible Peptides. *Journal of Chemical Theory and Computation*, 20(14):5829–5841, July 2024. ISSN 1549-9618. doi: 10.1021/acs.jctc.4c00502.

33 Yang Zhou, Rongfeng Zou, Guanglin Kuang, Bengt Långström, Christer Halldin, Hans Ågren, and Yaoquan Tu. Enhanced Sampling Simulations of Ligand Unbinding Kinetics Controlled by Protein Conformational Changes. *Journal of Chemical Information and Modeling*, 59(9): 3910–3918, September 2019. ISSN 1549-9596. doi: 10.1021/acs.jcim.9b00523.

34 Justin Spiriti and Chung F. Wong. Quantitative Prediction of Dissociation Rates of PYK2 Ligands Using Umbrella Sampling and Milestoning. *Journal of Chemical Theory and Computation*, 20(9):4029–4044, May 2024. ISSN 1549-9618. doi: 10.1021/acs.jctc.4c00192.

35 Samith Rathnayake, Brajesh Narayan, Ron Elber, and Chung F. Wong. Milestoning simulation of ligand dissociation from the glycogen synthase kinase 3. *Proteins: Structure, Function, and Bioinformatics*, 91(2):209–217, 2023. ISSN 1097-0134. doi: 10.1002/prot.26423.

36 Justin Spiriti, Frank Noé, and Chung F. Wong. Simulation of ligand dissociation kinetics from the protein kinase PYK2. *Journal of Computational Chemistry*, 43(28):1911–1922, 2022. ISSN 1096-987X. doi: 10.1002/jcc.26991.

37 Muhammad Jan Akhunzada, Hyun Jung Yoon, Indrajit Deb, Abdennour Braka, and Sangwook Wu. Bell-Evans model and steered molecular dynamics in uncovering the dissociation kinetics of ligands targeting G-protein-coupled receptors. *Scientific Reports*, 12(1):15972, September 2022. ISSN 2045-2322. doi: 10.1038/s41598-022-20065-2.

38 Abdennour Braka, Norbert Garnier, Pascal Bonnet, and Samia Aci-Sèche. Residence Time Prediction of Type 1 and 2 Kinase Inhibitors from Unbinding Simulations. *Journal of Chemical Information and Modeling*, 60(1):342–348, January 2020. ISSN 1549-9596. doi: 10.1021/acs.jcim.9b00497.

39 Shinji Iida and Tomoshi Kameda. Dissociation Rate Calculation via Constant-Force Steered Molecular Dynamics Simulation. *Journal of Chemical Information and Modeling*, 63(11):3369–3376, June 2023. ISSN 1549-9596. doi: 10.1021/acs.jcim.2c01529.

40 Kayla A. Croney and James McCarty. Exploring Product Release from Yeast Cytosine Deaminase with Metadynamics. *The Journal of Physical Chemistry B*, 128(13):3102–3112, April 2024. ISSN 1520-6106. doi: 10.1021/acs.jpcb.3c07972.

41 Karen Palacio-Rodriguez, Hadrien Vroylandt, Lukas S. Stelzl, Fabio Pietrucci, Gerhard Hummer, and Pilar Cossio. Transition Rates and Efficiency of Collective Variables from Time-Dependent Biased Simulations. *The Journal of Physical Chemistry Letters*, 13(32):7490–7496, August 2022. doi: 10.1021/acs.jpclett.2c01807.

42 Benjamin R Jagger, Anupam A Ojha, and Rommie E Amaro. Predicting ligand binding kinetics using a markovian milestoning with voronoi tessellations multiscale approach. *Journal of Chemical Theory and Computation*, 16(8):5348–5357, 2020.

43 Anupam Anand Ojha, Lane William Votapka, and Rommie Elizabeth Amaro. Qmrebind: incorporating quantum mechanical force field reparameterization at the ligand binding site for improved drug-target kinetics through milestoning simulations. *Chemical Science*, 14(45): 13159–13175, 2023.

44 Anupam Anand Ojha, Lane William Votapka, Gary Alexander Huber, Shang Gao, and Rommie Elizabeth Amaro. An introductory tutorial to the seekr2 (simulation enabled estimation of kinetic rates v. 2) multiscale milestoning software [article v1. 0]. *Living Journal of Computational Molecular Science*, 5(1):2359–2359, 2023.

45 Xiangyi Lu, Li Xiao, Luan Wang, and Douglas M Ruden. Hsp90 inhibitors and drug resistance in cancer: the potential benefits of combination therapies of hsp90 inhibitors and other anti-cancer drugs. *Biochemical pharmacology*, 83(8):995–1004, 2012.

46 Len Neckers, Brian Blagg, Timothy Haystead, Jane B Trepel, Luke Whitesell, and Didier Picard. Methods to validate hsp90 inhibitor specificity, to identify off-target effects, and to rethink approaches for further clinical development. *Cell Stress and Chaperones*, 23(4): 467–482, 2018.
